# Fusion-derived phospho-neoepitopes define a prioritized candidate neoantigen repertoire in MASLD-HCC

**DOI:** 10.64898/2026.08.26.747243

**Authors:** Li Na Zhao, Jesper B. Andersen

**Author notes:** **Lead author** Prof. Jesper. B. Andersen; Ole Maaløes Vej 5, Copenhagen N, DK-2200 Denmark. Correspondence may be addressed to either Professor Andersen or Dr. Lina Zhao.

## Abstract

**Background:** The rising burden of metabolic dysfunction-associated steatotic liver disease (MASLD)-associated hepatocellular carcinoma (HCC) underscores the need for innovative therapeutic strategies.

**Methods:** We integrated RNA-seq fusion detection, immunopeptidomics, and proteogenomics to systematically prioritize tumor-specific neoantigen candidates arising from gene fusions in MASLD-HCC.

**Results:** We elucidated a landscape of private, clonally expressed fusions, and identified a previously unrecognized class of predicted phosphorylated fusion-neoepitopes. Cross-tumor proteomic analysis revealed that these phospho-motifs are present across malignancies, providing a broader context for their biological relevance. Importantly, fusion-positive tumors display immunosuppressive microenvironments, highlighting the need for future therapeutic strategies that combine fusion-targeted immunotherapy with approaches that overcome T-cell dysfunction.

**Conclusions:** This study establishes a discovery pipeline and publicly available resource for fusion-derived phospho-neoepitopes in MASLD-HCC. The identified candidates provide a prioritized framework to guide and accelerate rigorous functional immunogenicity testing for future clinical validation.

**Highlights:**

- The landscape of MASLD-HCC fusions is dominated by predominantly private, with rare recurrent candidate genetic events
- Proteogenomic data supports high-confidence fusion-derived candidate neoepitopes
- Novel phosphorylated fusion-derived candidate neoepitopes are identified in MASLD-HCC
- Immunotherapy-treated HCC patients inform personalized fusion-derived candidate neoepitope discovery

**Graphical abstract:** 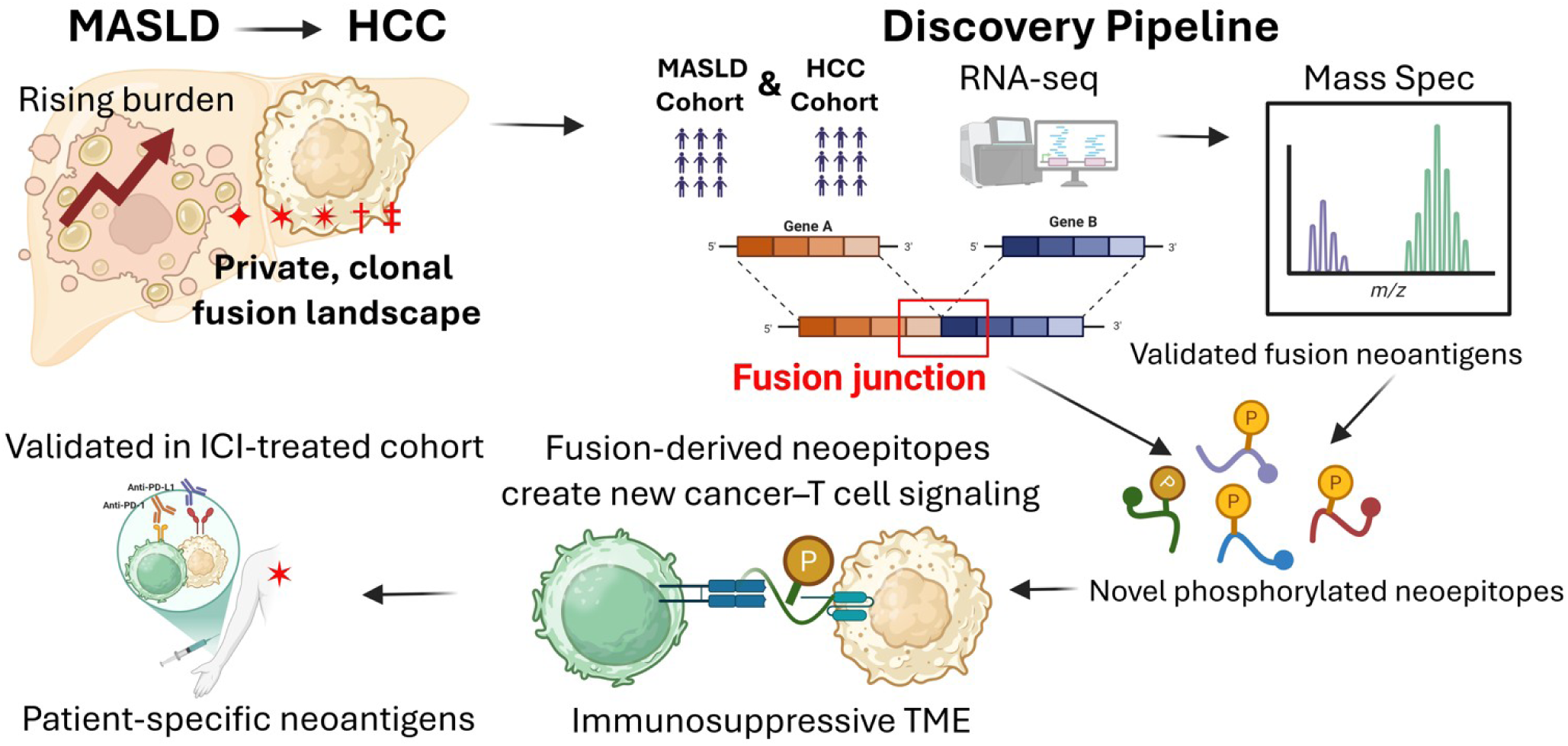

## INTRODUCTION

The global incidence of metabolic dysfunction-associated steatotic liver disease (MASLD) and MASLD-associated hepatocellular carcinoma (HCC) is rising at an alarming rate, positioning it as a major public health challenge, forecasting that MASLD alone will affect the global population by 25% in 2030 and contribute 40% to HCC[1]. Despite advances in immunotherapy, a significant proportion of MASLD-HCC patients exhibit poor response to immune checkpoint inhibition (ICI), often attributed to a non-inflamed, or immunologically cold, tumor microenvironment (TME)[2]. This clinical reality underscores the urgent need to identify novel, potent, and tumor-specific antigens capable of eliciting robust anti-tumor immunity[3,4].

Tumor-specific neoantigens arising from somatic alterations represent ideal drug targets due to their absence from healthy tissues, maximizing on-target tumor cell responses and reducing off-target toxicity[5,6]. While focus is placed on mutation-derived neoantigens, their clinical efficacy is limited by inter-and intratumor heterogeneity and often low immunogenic potency[7]. Contrary, gene fusions, resulting from chromosomal rearrangements, are compelling alternative sources of neoantigens[8]. Fusion events often generate entirely novel amino acid sequences specific at the fusion junction that are predominantly clonal and truncal events, making them resilient to tumor evolution and thus, ideal targets[9,10]. The potential for fusion-derived peptides to undergo post-translational modifications, such as phosphorylation, is largely unexplored and presents a potential for escalating the neoantigen repertoire[11,12]. Indeed, the fusion-derived neoantigen landscape in MASLD-HCC remains limited[13–15]. To overcome this restraint, we studied whether fusions are predominantly private events, with a subset of recurrent candidates, presenting candidate neoepitopes within the immunosuppressive MASLD-HCC TME. We performed an integrated genomics-to-proteogenomics analysis, mapping gene fusions in MASLD-HCCs, emphasizing proteogenomic evidence supporting candidate fusion-derived neoepitopes and their predicted immunogenicity. Importantly, we elucidated a novel class of phosphorylated fusion-derived epitopes, phosphopeptide neoantigens, in MASLD-HCC.

## METHOD

### Patient cohorts

Prospective patient samples were collected from two cohorts: (i) a MASLD cohort comprising 42 healthy controls, 66 individuals with MASLD, and 11 with Metabolic Dysfunction-Associated Steatohepatitis (MASH; n = 119 samples[16]); and (ii) a MASLD-HCC cohort, which included 47 patients with paired tumor and adjacent non-tumor tissue samples (n = 94 samples[17]).

### Ethic consideration

The MASLD study protocol conforms to the ethical guidelines of the 1975 Declaration of Helsinki and was approved by Etikprövningsmyndigheten (Dnr 2024-07917-01) in Sweden. MASLD-HCC was performed following individual patient consent, local institutional review board approvals (IORG0003254; IRB00003888), and assessed by the Committee on Health and Research Ethics for the Capital Region of Denmark for use of archival material ( no. H-4-2016-FSP, 17029679). All patient datasets were anonymized.

All other methods and data set accessions can be found in the online supplementary information.

## RESULT

### Landscape and biological characteristics of gene fusions in MASLD-HCC

To elucidate relevant gene fusions in MASLD-HCC, we utilized a genomics-to-proteogenomics analysis across multiple patient cohorts (**Fig. 1A**), unraveling a dominant genomically instable landscape characterized by accumulation of gene fusions that are prevalently private, clonal events, with a small subset of recurrent candidates. Analyzing liver tissue samples obtained from healthy controls to non-malignant MASLD and MASH cases (n = 119) and malignant MASLD-HCC (n = 47 patients, 94 tumor and normal paired samples) revealed a more than 25-fold increase in the burden of high-confidence fusions. Using stringent filtering criteria, multiple algorithms, and requiring that fusions are recurrent across tumor samples, we define in total 114 fusions present in 41 HCCs (6 patients with no unique detectable fusion events) compared to only 5 fusions in MASLD livers (in only 4 out of 119 samples) (**Fig. 1B**). This analysis showed a widespread ongoing process of chromosomal rearrangements in progressed MASLD-HCCs compared to the underlying MASLD livers, in which the parenchyma maintains a relative genomic stability. Though fusion events are heterogeneous across the cohort, with the majority unique to the individual patient, each fusion is present in the dominant proportion of tumor cells, indicating these events are generally clonal, early truncal events during the tumor evolution (**Fig. S1, S2**). Overall, recurrent gene fusions were identified across all tumor samples defining a core set of 23 events (**Fig. 1B-C, Table S1-2**). Among these fusions, the most prevalent (ADGRV1--TERT and CPS1--LANCL1-AS1) were found to be shared also across two independent cohorts, including our treatment naïve discovery cohort of MASLD-HCC and an external NCI-ICI treated validation cohort[18].

**Figure 1.**
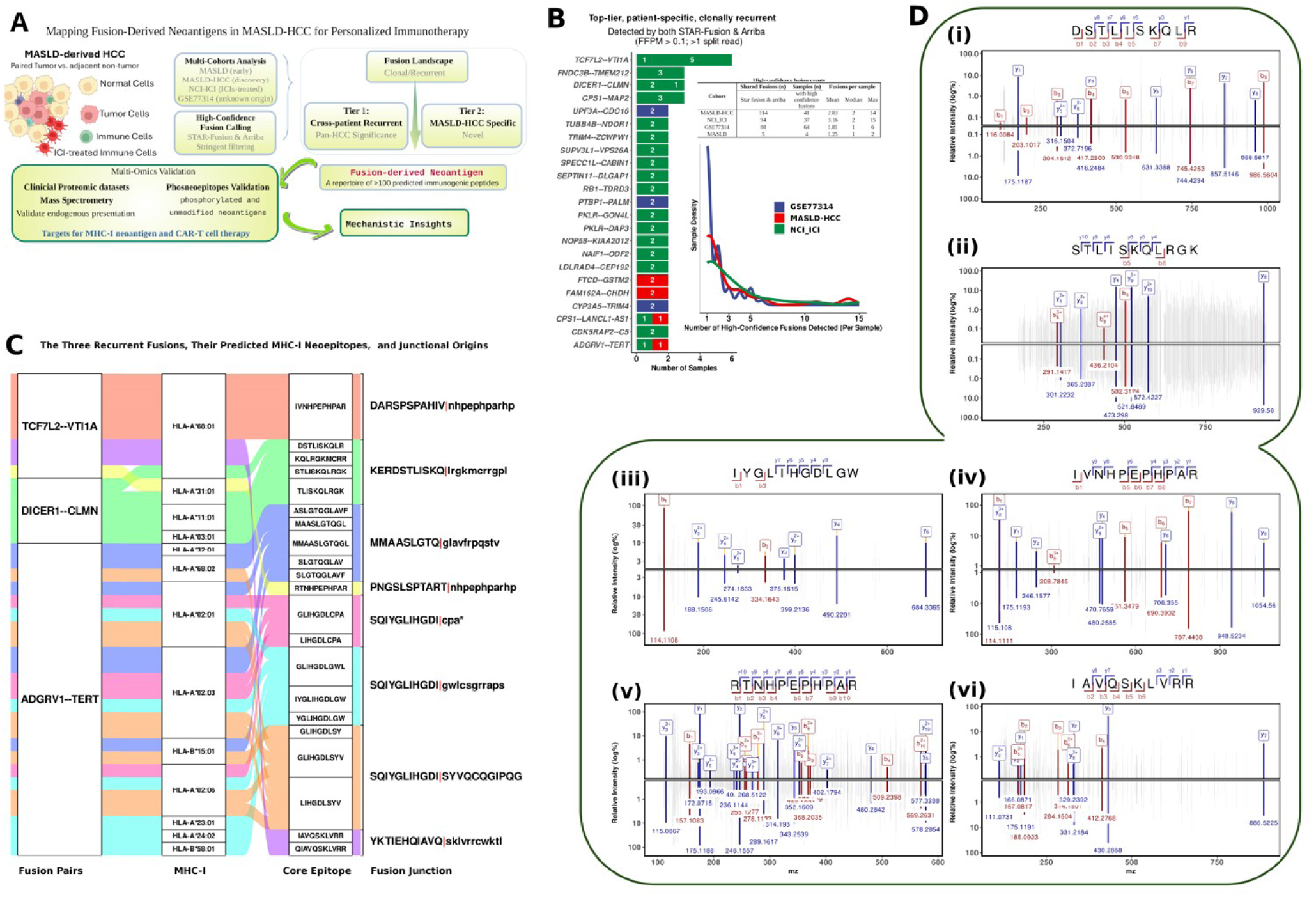
Integrated genomics-to-mechanism analysis uncovers oncogenic and immunogenic fusion drivers in MASLD-associated hepatocellular carcinoma. **(A)** An integrated genomics-to-mechanism workflow identifies actionable gene fusions in MASLD-HCC. **(B)** Distribution of high-confidence fusion events per sample across cohorts. Density plots show the number of fusion events detected per sample, restricted to fusions identified by both STAR-Fusion and Arriba (high-confidence shared fusions). **(C)** Mapping fusion-derived neoantigen presentation across MHC alleles and fusion junction. Alluvial diagram illustrating the relationships among fusion events, their predicted neoantigen peptides, corresponding HLA class I alleles, and the fusion-junction peptide segments. For each fusion, only the 11 amino acids flanking either side of the junction (left₁₁|right₁₁) were analyzed, with the junction site indicated by a red vertical bar (“|”). **(D)** Proteogenomic support for fusion-derived candidate neoepitopes by MS/MS. DICER1-CLMN: (i) Peptide *DSTLISKQLR* from the CPTAC HNSCC proteome (PDC000221; spectrum 13CPTAC_HNSCC_W_JHU_20190731_LUMOS_f02:32697:2); (ii) Peptide *STLISKQLRGK* from the CCLE proteome (MSV000085836; spectrum: g00804_Prot_25_09:33864:2); (iii) ADGRV1--TERT: Peptide *IYGLIHGDLGW* from the gastric cancer proteome (PDC000214; spectrum: FN04_N55T56_180min_10ug_C2_032813:8779:3); TCF7L2--VTI1A: (iv) Phosphopeptide *IVNHPEPHPAR* from the ovarian cancer phosphoproteome (PDC000115; spectrum: TCGA_25-1628_13-1494_24-1104_117C_P_PNNL_B1S4_f08:9055); (v) Peptide *RTNHPEPHPAR* from the pancreatic ductal adenocarcinoma (PDA) proteome (PDC000270; spectrum: 13CPTAC_PDA_W_JHU_20191219_LUMOS_f12:8661:3) ; (vi) Peptide *IAVQSKLVRR* from the breast cancer proteome (PDC000120; spectrum: 10CPTAC_Bcprospective_W_BI_20170224_BL_f20:46228:2). Matched b-ions and y-ions shown in red and blue, respectively. All spectra show high-confidence prediction (PSM scores 15.6 - 24.2, mass tolerance = 0.06 Da). Peptide-spectrum matches were validated by PepQuery, confirming they pass all criteria for novel peptide identification.

To refine therapeutically actionable targets from this complex fusion landscape, we implemented a multi-tiered prioritization framework that integrated the requirement for a fusion to be recurrent, protein coding, and present a minimal likelihood for off-tumor cross-reactivity. As such, prioritizing fusions predicted to generate tumor-specific sequences with minimal expression or homology in the normal liver parenchyma.

We prioritized shared fusions across several cohorts, adding a filter focusing on the biological recurrence to emphasize its potential in designing therapies. This limited our previously defined 23 recurrent fusions (**Fig. 1B**) to three clonal, tumor-exclusive events, including TCF7L2--VTI1A, DICER1--CLMN, and ADGRV1--TERT (**Fig. 1C, Table S1**). Among the selected fusions, ADGRV1--TERT presents the only protein-coding fusion, positioning it as a high-priority MHC-I candidate neoantigen.

These prioritization steps critically informed our *in silico* safety screen, highlighting core epitopes derived from the selected fusions using large-scale global tissue proteomes curated from healthy donor tissue datasets in PepQuery[19–21]. This analysis revealed in total 8 significant peptides, with 6 of these epitopes being confidently tumor-specific. The remaining two peptides (MMAASLGTQGL and SLGTQGLAVF) were detected in deep proteomes from healthy human tissues, suggesting they may present potential off-tumor risks to the normal liver though they are on-target (**Fig. 1D**). Thus, we deprioritized these peptide sequences and limited our selection to the remaining 6 junctional tumor-specific peptides, presenting high-confident candidates for further evaluation.

Given that most fusions are private, patient-specific events, we applied an integrative strategy combining multiple fusion-calling algorithms to comprehensively characterize the fusion landscape in MASLD-HCC. This approach identified a repertoire of 55 shared and protein-coding fusions across the MASLD-HCC cohort (**Table S2**). Most of these fusion events are novel, revealing a distinct fusion landscape in MASLD-HCC characterized by marked heterogeneity and a predominance of non-recurrent structural rearrangements, rather than canonical, highly recurrent oncogenic fusions (**Fig. 2A**). Although the overwhelming majority of fusion events were private and patient-specific, a small subset of recurrent fusions, including ADGRV1–TERT, was identified across independent cohorts and prioritized for further analysis. Notably, most shared protein-coding fusions exclusively utilize reference splice junctions, indicating that these events are predominantly driven by underlying genomic rearrangements that juxtapose intact exons, rather than by aberrant or non-canonical splicing events (**Table S2**). This observation is consistent with prior large-scale analyses of TCGA-LIHC and independent HCC cohorts, which similarly demonstrated that the HCC fusion landscape is dominated by heterogeneous and largely non-recurrent fusion events, with only a minority of recurrent driver-like fusions shared across tumors[22–26]. Importantly, these studies also found that most fusion transcripts involve canonical exon boundaries and are likely generated by structural rearrangements rather than widespread aberrant splicing, supporting our interpretation that structural variation is a principal mechanism underlying fusion transcript formation in HCC[27,28]. Moreover, among the 55 shared fusions, a subset (20/55 fusion events) overlaps with known oncogenes in the fusion-gene database (FusionGDB2.0), supporting their potential causal pathological roles (**Fig. S2ab, Table S2**).

**Figure 2.**
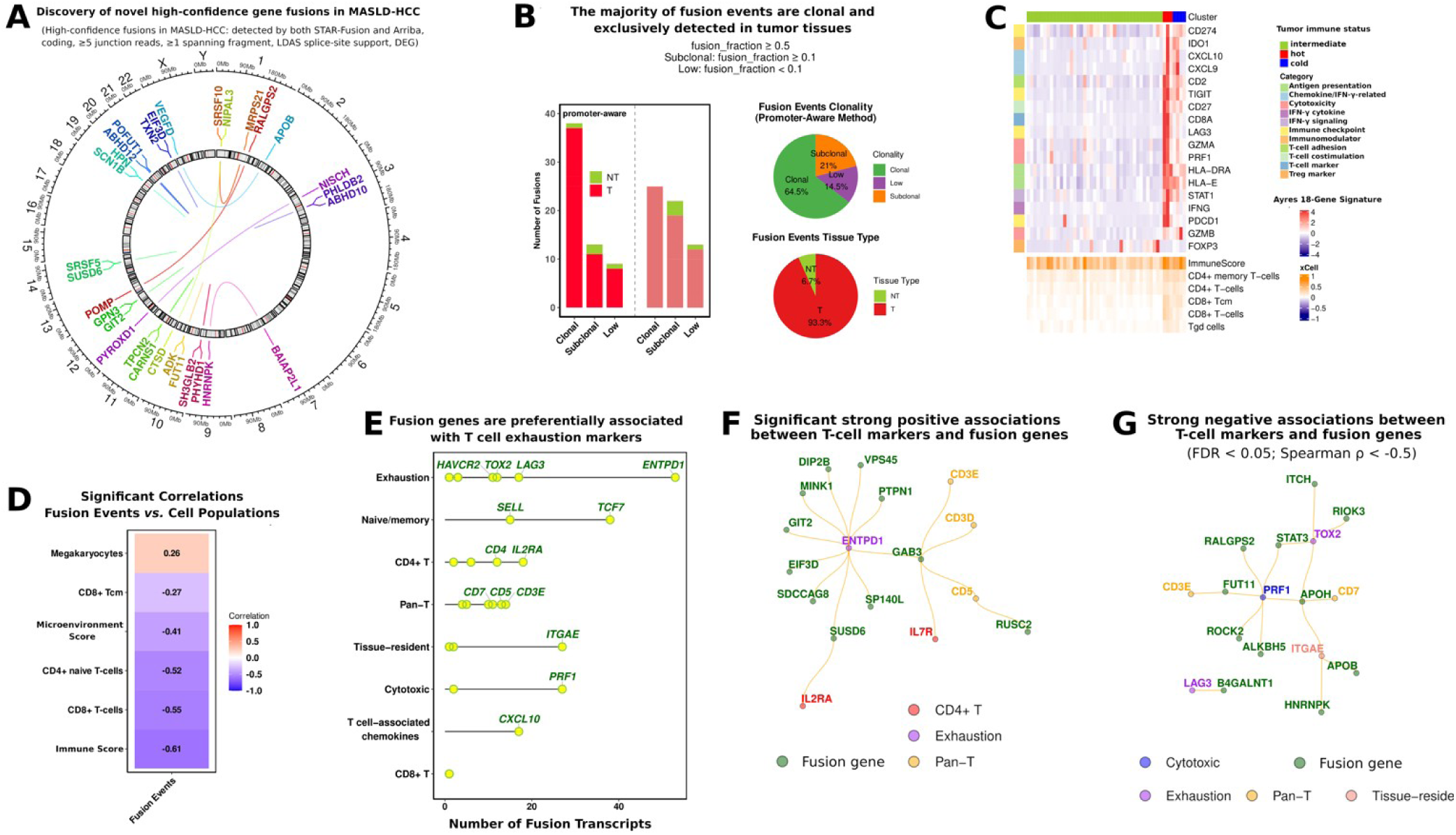
Gene fusions and immune microenvironment in MASLD-HCC. (**A**) Circos visualization of novel high-confidence gene fusions in MASLD-HCC cohort. The plot shows coding, differentially expressed gene fusions identified in MASLD-HCC cohort. Links connect chromosomal loci of fusion partners, with line thickness scaled to the number of supporting samples. Gene labels highlight fusion partners, and color-coding distinguishes unique fusion events. Only high-confidence fusions (≥5 junction reads, ≥1 spanning fragment, and splice-support criteria) with evidence of high junction read counts are shown. (**B**) Fusion event clonality. Stacked bar plots show the proportion of clonal, subclonal, and low-fraction fusions across samples, grouped by promoter-specific clonality (based on the promoter-bearing fusion partner) and overall clonality. (**C**) Immune landscape stratification of MASLD-HCC tumors reveals distinct microenvironmental patterns. Top: Unsupervised clustering of tumors based on the Ayres 18-gene T cell exhaustion signature, showing expression levels across samples. Bottom: Corresponding xCell enrichment scores for key immune and stromal cell populations. Samples are consistently ordered across both heatmaps and colored by immunologically cold (blue), intermediate (yellow/green), and immunologically hot (red). (**D**) Significant correlations between fusion events and immune cell populations in MASLD-HCC. (**E**) Lollipop plot showing the number of fusion genes significantly correlated with T cell markers (|ρ| > 0.4, FDR-adjusted < 0.05). Labels indicate T cell markers associated with more than 10 fusion transcripts. (**F**) Correlation networks between T cell transcriptional programs and fusion gene expression. Strong positive correlations (Spearman ρ > 0.6, FDR-adjusted p < 0.05) indicate genes co-expressed with T cell exhaustion markers ENTPD1 and CD4⁺ T cell markers IL7R and IL2RA (CD25). (**G**) Significant negative correlations (Spearman ρ ← 0.5, FDR-adjusted p < 0.05) highlight genes inversely associated with cytotoxic, exhausted, and tissue-resident T cell markers.

To prioritize biologically relevant candidates, we next filtered this set based on differential expression between tumor and matched non-tumor liver tissues, yielding 37 tumor-enriched fusions. Of these, 23 fusions are in-frame events with putative protein-coding function, including 5 fusions with known oncogenic roles and 16 novel predicted drivers (**Table S2**). Moreover, these 23 tumor-specific fusions are established by predominantly in-frame breakpoints occurring at protein-coding sequences or splice-site junctions, thereby preserving functional domains critical for oncogene function such as constitutive kinase activation as for the fibroblast growth factor receptor 2 and arginyltransferase 1 (FGFR2--ATE1) fusion event (**Fig. S2c-e**).

Next, to determine the evolutionary timing of these events, we assessed the clonality of high-confidence fusion transcripts across tumor samples. Fusion events were found to be predominantly tumor-specific and clonal, consistent with early truncal alterations that are maintained during tumor progression (**Fig. 2B**). In contrast, matched tumor-adjacent liver tissues rarely harbored detectable fusions, but when present, these events are classified as sub-clonal, occurring at low allele frequencies, which suggest that they are late-arising and likely represent passenger alterations rather than functionally relevant drivers.

Having established that most fusions in the tumors are of early and clonal origin, we next examined their functional consequences on the transcriptome. As such, a subset of fusion events is associated with pronounced transcriptional rewiring, highlighting diverse mechanisms by which structural rearrangements alter gene regulation. For instance, the recurring ADGRV1--TERT fusion results in suppression of both wild-type partner genes, with the fusion becoming the dominant expressed transcript (**Fig. S3ab**). In contrast, the LAMA3--RIOK3 fusion exemplifies a situation with promoter hijacking, whereby the chimeric transcript is expressed at a level 27-fold higher than the native *RIOK3* gene. Besides, we also identified evidence of concurrent two-hit inactivation of tumor suppressor genes, including the RERE--PLCH2 fusion, which results in complete loss of the wild-type *RERE* expression, consistent with a dual oncogenic mechanism involving both structural disruption and transcriptional silencing. Together, these findings demonstrate that fusion events can exert potent cis-acting effects on gene expression, including transcript dominance, promoter reassignment, and tumor suppressor inactivation. It should be noted, there is a significant difference in the sensitivity of detection between sequencing methods. Importantly, most fusion transcripts were captured by snRNA-seq (**Fig. S3c**; GSE189175), whereas only a few genes (such as, Apolipoprotein B (*APOB*), Transferrin (*TF*), Cathepsin D (*CTSD*)) were detected across several scRNA-seq datasets (GSE149614, GSE189903). This pattern, coupled with the generally higher detection limit of the 5’ fusion partner in snRNA-seq (**Fig. S3d**), indicates that a significant proportion of chimeric transcripts are pre-mRNAs or nuclear-retained RNA species. Moreover, the genomic breakpoints of these fusions predominantly occur within introns, or at exon-intron junctions, and that the primary fusion transcript is subject to nuclear processing before cytoplasmic export.

Taken together, the clonal nature, the potent cis-acting effects on gene expression, and the nuclear localization of these transcripts, demonstrate that these fusion events originate from structural variations in the tumor genomes.

### Fusion burden is associated with the immunosuppressive tumor microenvironment

Analysis of the treatment-naïve MASLD-HCC tumor microenvironment revealed that, compared to tumor-adjacent tissues, tumors exhibited endothelial enrichment and depletion of fibroblasts, platelets, and multiple immune components (**Fig. S4a**). In this context, fusion events were associated with concurrent vascular expansion and immune suppression. This fusion-burden positively correlated with endothelial/vascular signatures, while showing a negative correlation with CD8⁺ T cell infiltration and immune activity (assessed by the Ayres immune gene signature[29]; **Fig. S4b**), suggesting that fusion-positive tumors may promote an immunologically cold tumor.

To examine this observation, we obtained data from the NCI-ICI cohort, showing that following immunotherapy this relationship still persists regardless that tumors maintain reduced immune scores compared to the tumor-adjacent liver (**Fig. S4c**). Following immunotherapy, fusion events are positively correlated with CD4⁺ and memory T cell populations, contrasting the broadly immunosuppressed phenotype observed in treatment-naïve tumors (**Fig. S4d**). To systematically classify the immune contexture, we stratified tumors into either immunologically cold, intermediate, or hot using Ayres and xCELL cellular deconvolution and enrichment scores. Most tumors are associated with the intermediate category, presenting a distinct distribution of various T cell populations (CD8⁺, CD4⁺, CD4⁺ memory, CD8⁺ central memory, and γ/δ T cells) across the three clusters (**Fig. 2C, S4e**). Overall, the fusion-burden did not correlate with the immune classes but showed significant negative associations with prediction of the ImmuneScore, CD8⁺ T cells, and naïve CD4⁺ T cells, supporting a role for fusion-driven alterations in promoting an immune-excluded tumor phenotype (**Fig. 2D, S4f**).

Utilizing single-cell transcriptomics it has been shown that tumor-reactive CD8⁺ T cells in liver cancer[30] and in other solid tumors[31] primarily are found in exhausted, tissue-resident clusters characterized by high (PD-1, TIM-3, LAG3, TIGIT, TOX, CD39, CD69, CD103, CXCR6) and low TCF7 marker expression. In our analysis, fusion genes displayed a significantly stronger and more frequent association with immune exhaustion markers (ENTPD1) and CD4⁺ T cell markers (IL7R and IL2RA/CD25)) than to other markers, suggesting that fusion-positive tumors may be associated with T cell dysfunction and CD4⁺ T cell activity (**Fig. 2E-F**). Conversely, significant anticorrelations were observed with cytotoxic and tissue-resident markers (PRF1, LAG3, and TOX2), consistent with reduced cytolytic activity in fusion-enriched tumors (**Fig. 2G**).

### Proteogenomic integration prioritizes fusion-derived candidate neoepitopes

Next, we sought to determine the immunogenic potential of the defined set of high-confidence fusions. To achieve this, we developed a proteogenomic integration moving beyond *in silico* prediction by providing orthogonal mass spectrometry evidence for the endogenous expression and proteolytic processing of fusion-derived peptides, thereby prioritizing high-confidence candidate neoepitopes for downstream functional validation. We used the Immune Epitope Database (IEDB) next-generation platform to analyze peptides spanning the fusion junctions of shared, in-frame fusions against a reference panel of 27 HLA class I alleles by integrating metrics for MHC-I binding, antigen processing, and potential for immunogenicity, identifying 70 potential immunogenic peptides (**Fig. 3A**). To confirm which of the 70 predicted neoantigens are endogenously detected on the cell surface, we screened these peptides against a compendium of 48 publicly available proteomic datasets (>1 million MS/MS spectra)[32], providing significant evidence for 12 prioritized fusion-derived peptides (12/70 peptides) (**Table S2**). Subsequently, we applied a set of stringent quality filters, excluding first two peptides based on low MS/MS intensity and weak identification scores. Lastly, we analyzed the final 10 peptides against proteomes obtained from healthy donor tissues, removing four additional peptides, leaving 6 peptides to be prioritized as high-confidence tumor-restricted candidate peptides ( **Fig. 3B, S5a, Fig. S6a-c**).

**Figure 3.**
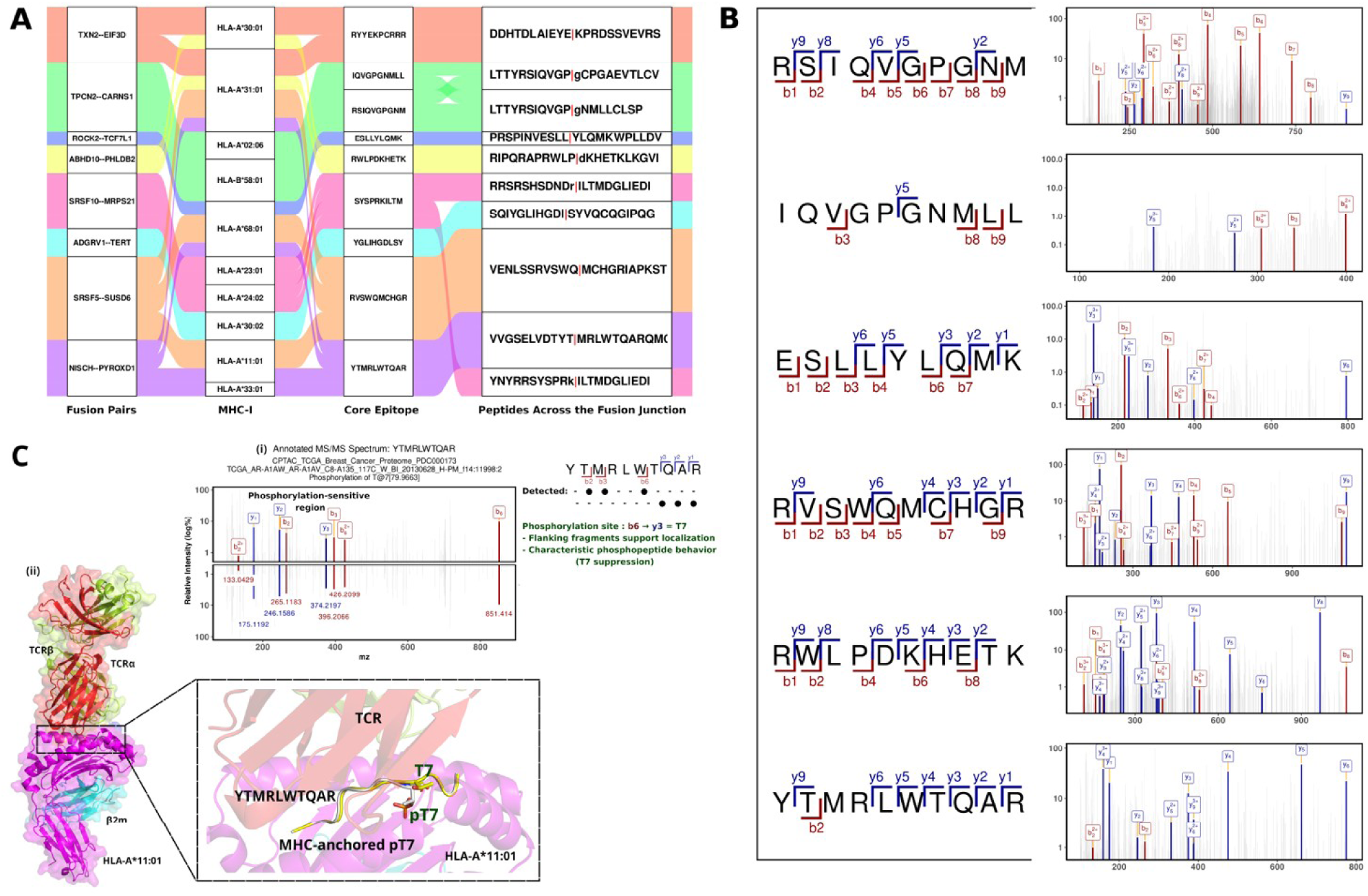
Discovery and characterization of fusion-derived and phosphorylated neoantigens in MASLD-HCC. (**A**) Alluvial representation of top fusion-derived immunogenic peptides that are detected in the clinical proteomic datasets. (**B**) Detection of a phosphorylated neoepitope in the MASLD-HCC cohort. (**C**) Detection and structural modeling of the phosphorylated YTMRLWTQAR neoepitope. (i) Annotated MS/MS spectrum confirming the detection of the phosphorylated (pT7) form of the YTMRLWTQAR peptide in clinical proteomes. Key b-and y-ions are labeled. (ii) Structural model of the neoepitope within the MHC binding groove. Superposition of the unphosphorylated (yellow) and phosphorylated (white) peptides *raises the possibility of an altered pMHC conformation, though further studies (crystallography or binding assays) are required to confirm this*.

### Fusion-derived neoantigens exhibit posttranslational phosphorylation

Importantly, we have identified phosphorylated candidates peptides derived directly from gene-fusion products. These phospho-neopeptides represent a dual anomaly, by arising both from genomic rearrangement and following a cancer-associated posttranslational modification (PTM). Overall, we identified 15 high-confidence tumor-specific phospho-neopeptides, of which 10 sites were confirmed in clinical cancer proteomes[33] and by the Clinical Proteomic Tumor Analysis Consortium **(Table S2, Fig. S7a**), including the peptides TTPYGGVSRR (from SH3GLB2--PHYHD1) and ESLLYLQMKW (from ROCK2--TCF7L1). Structural analysis revealed phosphorylation at critical positions, including T cell receptor (TCR)-facing residues (ESLL**pY**LQMKW and STLI**pS**KQLRGK) and MHC anchor residues (**pTpT**PYGGVSRR), which are predicted to substantially alter the immune recognition. Besides, we detected the fusion peptide YTMRLWTQAR that is derived from the fusion NISCH--PYROXD1 in both unphosphorylated state and with phosphorylation on threonine 7 (YTMRLW**pT**QAR), demonstrating a wider posttranslational plasticity in the overall antigen presentation (**Fig. 3B-C**). Moreover, the core motif LWTQAR aligned with the peptide derived from NISCH--PYROXD1 also constitutes a potential antigenic hotspot that overlaps with the known MHC-I ligand R**LWTQAR**QMGW (**Fig S7b;** IEDB ID:21231898)[34–37]. Additionally, this is significantly supported by structural modelling and the documented presence of methionine-oxidized variants of this ligand that indicates this region tolerates multiple PTMs (**Fig. S7c**).

A structural mining survey of all available antibody-antigen, TCR-pMHC, and MHC-ligand complexes in the Protein Data Bank revealed no crystal structures containing phospho-threonine (pT) moieties. In contrast, all crystallized phospho-serine (pS) modifications (n = 15) are oriented toward the TCR-facing surface, positioning them on the exposed interface of the antigen-presenting cell and enabling potential engagement with the T-cell receptor (**Fig. S7b**). This observation suggests that pT residues may similarly induce conformational changes that alter peptide-MHC topology and modulate TCR recognition. Similar, pT-specific modeling indicates that pT modifications likewise may induce conformational adjustments capable of modulating antigen presentation (**Fig. 3D**).

To determine whether such phosphorylation events can confer tumor specificity to otherwise shared peptide backbones, we next examined peptides detected across both healthy and tumor tissues. We identified a clear distinction in phosphorylation-dependent specificity for the peptide ISMLSSLGYI derived from fusion BRD9--ABCA3. In its unphosphorylated form (oM3; **Table S2**) this peptide was detected in healthy tissues, while the phosphorylated variant (pY9+oM3; **Table S2; Fig. S7a**) was shown only in tumors. This pattern indicates the possibility for phosphorylation to convert a peptide backbone that otherwise is shared with normal tissues into instead a tumor-specific epitope, underscoring the importance of considering PTMs when designing immunotherapies to minimize possible adverse effects. Additionally, DISMLSSLGY (also derived from BRD9--ABCA3) was found in two modified forms (pS3+pS6 and oM4+pS3) only in tumors, further highlighting PTM/phosphorylation as a key determinant of tumor-restricted antigen presentation.

Importantly, these observations are not unique to MASLD-HCC, with similar data shown in the *Proteogenomics of Gastric Cancer Glycoproteome* (PDC000216)[38], in which the peptide pSpYpSPRKILTM was detected with multiple phosphorylation sites and oxidized amino acid at M10. The repetition of modified peptides across distinct tumor types reinforces the concept that fusion-derived phospho-peptides constitute a broad and previously unrecognized source of tumor-specific neoantigens.

### Personalized prioritization of candidate neoepitopes

Our proteogenomic pipeline provides a comprehensive resource of fusion-derived candidate neoepitopes. To evaluate their clinical relevance at the individual level, we performed clinical-grade HLA typing. This analysis revealed a key limitation of broad screening approaches as many patient-specific HLA alleles (48 of 71; **Table S2**) were not represented in the predefined 27-allele human MHC class I panel commonly used in epitope prediction tools from the IEDB. As a result, the *in silico*-derived candidates were substantially reduced to five high-confidence, patient-matched neoepitopes corresponding to five different fusions (SRSF10--MRPS21; TPCN2--CARNS1; SRSF5--SUSD6; SH3GLB2--PHYHD1; ADGRV1--TERT), highlighting that, while large-scale discovery pipelines are valuable for defining a landscape of shared (“public”) candidates, such as ADGRV1–TERT, clinical translation ultimately requires a fully personalized strategy grounded in a patient’s own unique HLA haplotype and private fusion repertoire.

We next prioritized the ADGRV1–TERT fusion as a leading candidate based on its favorable structural and immunological properties. The fusion generates a novel junctional epitope (YGLIHGDLSY) while retaining the ADGRV1 extracellular domain, enabling potential targeting through both MHC-I-restricted vaccines and CAR-T approaches (**Fig. 4A-D, S3A**). Chromatin profiling indicated TERT promoter activation consistent with promoter hijacking (**Fig. 4E**), alongside suppression of the wild-type TERT transcript (**Fig. S4g**), supporting its role as a dominant oncogenic driver. Its tumor-restricted expression further supports therapeutic targeting of the fusion junction epitope to achieve selective tumor recognition. Notably, no recurrent phosphorylation events were detected within this epitope, suggesting stable presentation of an unmodified neoantigen.

**Figure 4.**
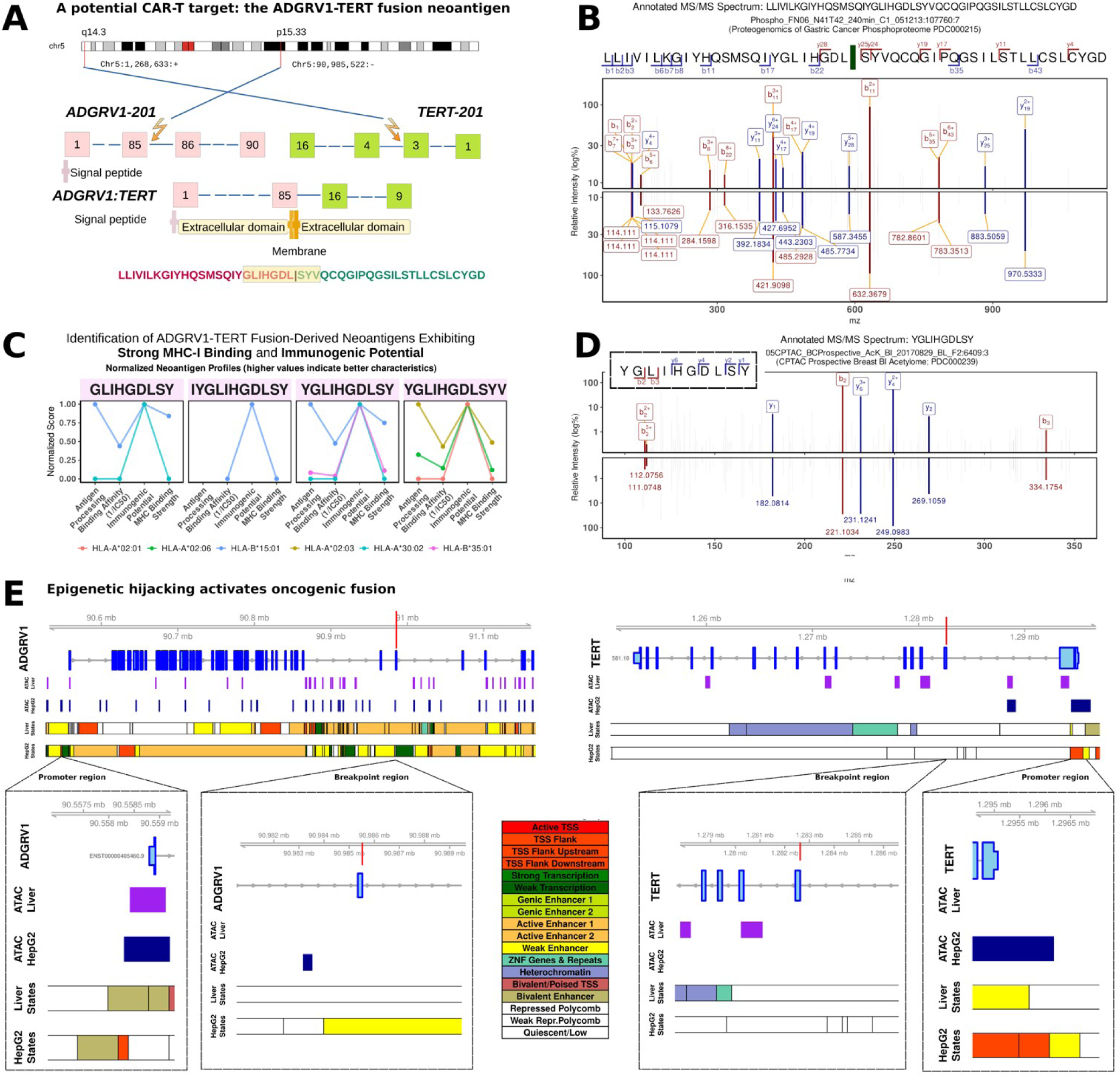
ADGRV1–TERT: A recurrent fusion-derived candidate neoantigen representing a rare exception within a predominantly private fusion landscape. (**A**) Schematic representation of the ADGRV1--TERT fusion. The fusion arises from an intrachromosomal inversion on chromosome 5, generating an in-frame transcript that joins the 5′ region of ADGRV1 to the 3′ region of TERT. The resulting chimeric protein retains the extracellular domains of ADGRV1 fused to TERT. Notably, wild-type TERT is largely silenced, making the fusion the dominant functional form. Multiple breakpoints in TERT produce several fusion transcripts, all in-frame. Notably, one fusion transcript retains an extracellular fusion junction, highlighting its potential as a candidate target for antibody-or CAR-based therapeutic strategies, pending experimental validation. (**B**) Tandem mass spectrum confirming the confident detection of the ADGRV1--TERT fusion peptide (LLIVILKGIYHQSMSQIYGLIHGDLSYVQCQGIPQGSILSTLLCSLCYGD) across multiple cancer proteomes. The fusion neoantigen candidate was strictly identified in clear cell renal cell carcinoma (PDC000127), glioblastoma (PDC000204) , lung adenocarcinoma (PDC000153, PDC000224), endometrial carcinoma (PDC000126), and gastric cancer (PDC000215) datasets, demonstrating pan-cancer expression. Key fragment ions are annotated, supporting the peptide sequence across diverse modification states including oxidation, carbamidomethylation, and methylation. (**C**) Identification of ADGRV1--TERT fusion-derived neoantigens with strong MHC-I binding and immunogenic potential. This analysis highlights candidate peptides spanning the ADGRV1-TERT fusion junction containing the “DLSY” motif. Four high-scoring variants, GLIHGDLSY, YGLIHGDLSY, YGLIHGDLSYV, and IYGLIHGDLSY, exhibited favorable MHC-I binding affinities (IC50 < 500 nM), low percentile ranks (<2%), and positive immunogenicity scores. Normalized neoantigen profiles are shown for each peptide, illustrating consistent trends across MHC alleles (higher values indicate more immunogenic characteristics). (**D**) Annotated MS/MS spectra confirming the immunogenic peptide YGLIHGDLSY derived from the ADGRV1–TERT fusion. Representative tandem mass spectra illustrate fragment ion annotation for the immunogenic peptide YGLIHGDLSY, derived from the ADGRV1--TERT fusion, detected in the DC000239 dataset. The upper panel displays annotated *b* and *y* fragment ions labeled with ion types, while the lower panel shows the corresponding fragment ion *m/z* values. Matched ions are colored in dark red (*b*-ions) and dark blue (*y*-ions), whereas unmatched peaks appear in gray. The intensity axis is presented on a logarithmic scale, and all annotated ions were manually verified. These spectra provide confident evidence supporting the presence of the ADGRV1-TERT fusion-derived immunogenic peptide in clinical tumor proteomic data. (**E**) Chromatin accessibility and state annotations at the ADGRV1--TERT fusion locus. Genome browser view of the ADGRV1 (left) and TERT (right) loci, showing gene models, ATAC-seq signal in healthy liver and HepG2, and ChromHMM state annotations. Vertical lines indicate the genomic breakpoints.

Importantly, comprehensive interrogation of PTM immunopeptidomes did not reveal recurrent phosphorylation events within the core ADGRV1-TERT junction epitope, suggesting that its immune recognition is predominantly driven by a constant presentation of an unmodified, high-affinity neoepitope.

Moreover, to further explore how PTMs may diversify fusion-derived antigenicity we characterized the SH3GLB2--PHYHD1 fusion, which is among the events generating MS-detected phospho-neopeptides (**Fig. 5A, S7a**).While PHYHD1 exhibits reduced chromatin accessibility in HepG2 cells and consistently is downregulated in HCC tumors (**Fig. 5B**; log_2_FC = -1.2, TCGA), the fusion places C-terminal *PHYHD1* exons under control of the highly active *SH3GLB2* promoter (log_2_FC = 0.6), which is upregulated in HCC. This junction disrupts the pyridine nucleotide-disulfide oxidoreductase domain of PHYHD1, generating a subpopulation of tumor cells that express a truncated, potentially non-functional PHYHD1 fragment. Rather than preserving canonical redox function, this truncated isoform may be non-functional or may perturb endogenous PHYHD1 activity while simultaneously creating novel antigenic determinants that reshape cellular fitness and immune recognition within the TME. Epitope prediction under HLA-B*07:02 identified three strong MHC binders derived from the fusion junction (**Fig. 5A**). Strikingly, high-confidence phosphopeptide-spectrum matches pinpointed the exact fusion junction-derived peptide and revealed extensive PTMs, impacting the core MHC-I epitope (TPYGGVpSRRM) where pS faces the TCR predicting to disrupt MHC-I binding and alter T cell recognition (**Fig. 5C**). This highlights a novel mechanism of immune regulation for fusion-derived neoantigens and underscores the critical need for phospho-immunopeptidomic validation.

**Figure 5.**
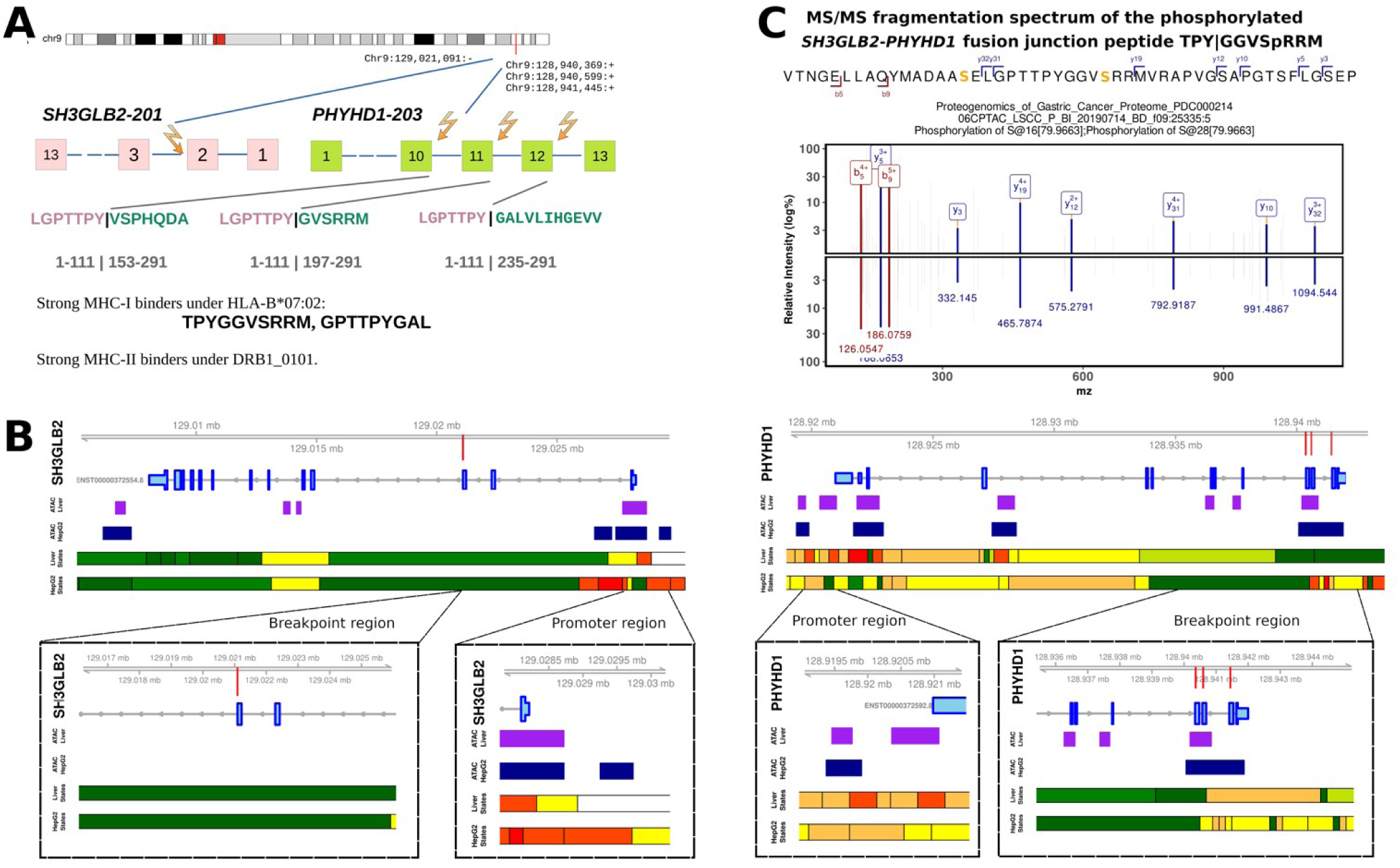
Genomic and proteomic characterization of the SH3GLB2--PHYHD1 gene fusion in hepatocellular carcinoma. (**A**) Schematic representation of the SH3GLB2--PHYHD1 fusion. This fusion results from an intrachromosomal inversion on chromosome 9, generating an in-frame transcript that joins the 5′ region of SH3GLB2 to the 3′ region of PHYHD1. The chimeric protein retains the N-terminal BAR domain of SH3GLB2, fused to a partial Phytanoyl-CoA dioxygenase domain of PHYHD1. Multiple breakpoints in PHYHD1 give rise to several fusion transcripts. (**B**) Chromatin accessibility and state annotations at the SH3GLB2--PHYHD1 fusion locus. Genome browser tracks show (left) SH3GLB2 and (right) PHYHD1 loci with gene models, ATAC-seq signals in healthy liver and HepG2 cells, and ChromHMM state annotations. Vertical lines indicate fusion breakpoints. ( **C**) Annotated MS/MS spectrum of the fusion-spanning peptide derived from SH3GLB2--PHYHD1 detected in a clinical cancer proteomic dataset. This peptide spans the fusion junction and serves as proteogenomic evidence for endogenous expression of the fusion product. The spectrum was acquired from PDC000232 with precursor m/z 1025.87. The peptide contains phosphorylation modifications at S16 and S28 (highlighted in orange). Matched b-ions (red) and y-ions (blue) are labeled, with 9 high-confidence fragment matches identified using a 0.06 Da mass tolerance. Despite sparse fragmentation characteristic of large phosphopeptides, the detected fragments provide supporting evidence for this fusion event.

To go beyond the fusion products themselves, we integrated proteomic and transcriptomic data within a 10kb window of the breakpoints and identified genes with significantly altered expression levels between tumor and tumor-adjacent tissues (**Fig. 6A; Table S2**). Predominantly these are known HCC-associated genes with roles in angiogenesis, metabolic reprogramming, and cancer. The proximity of these alterations to fusion breakpoints suggests that fusions directly perturb local gene regulation through enhancer hijacking, promoter disruption, or chromatin remodeling, positioning them as potent *cis*-regulatory drivers of the tumor development and maintenance. Based on criteria including in-frame fusion architecture, preservation of key functional domains, and precise exon–exon breakpoints, features that together suggest a high probability of generating stable, functional chimeric proteins, we prioritized three specific in-frame fusions for mechanistic investigation (ROCK2--TCF7L1, HPN--SCN1B, and GIT2--GPN3). Specifically, ROCK2--TCF7L1 retains the N-terminal domain of ROCK2 linked to the high-mobility group (HMG) box of TCF7L1 (**Fig. 6B**), suggesting a neomorphic role aberrantly modulating Wnt/β-catenin-driven transcription. The HPN--SCN1B fusion inserts a partial serine protease domain of the *HPN* gene onto the immunoglobulin-like domain of SCN1B (**Fig. 6C**); while its catalytic activity is likely abolished, it may function as a dysregulated cell-surface receptor that may disrupt normal extracellular signaling and adhesion. Finally, the GIT2--GPN3 fusion places the essential GPN3 GTPase under the control of the GIT2 promoter (**Fig. 6D**), and the resulting protein lacks residues critical for ATP binding (**Fig. 6E**), which is predicted to cripple its role in RNA Polymerase II assembly and thus, disrupt global transcription.

**Figure 6.**
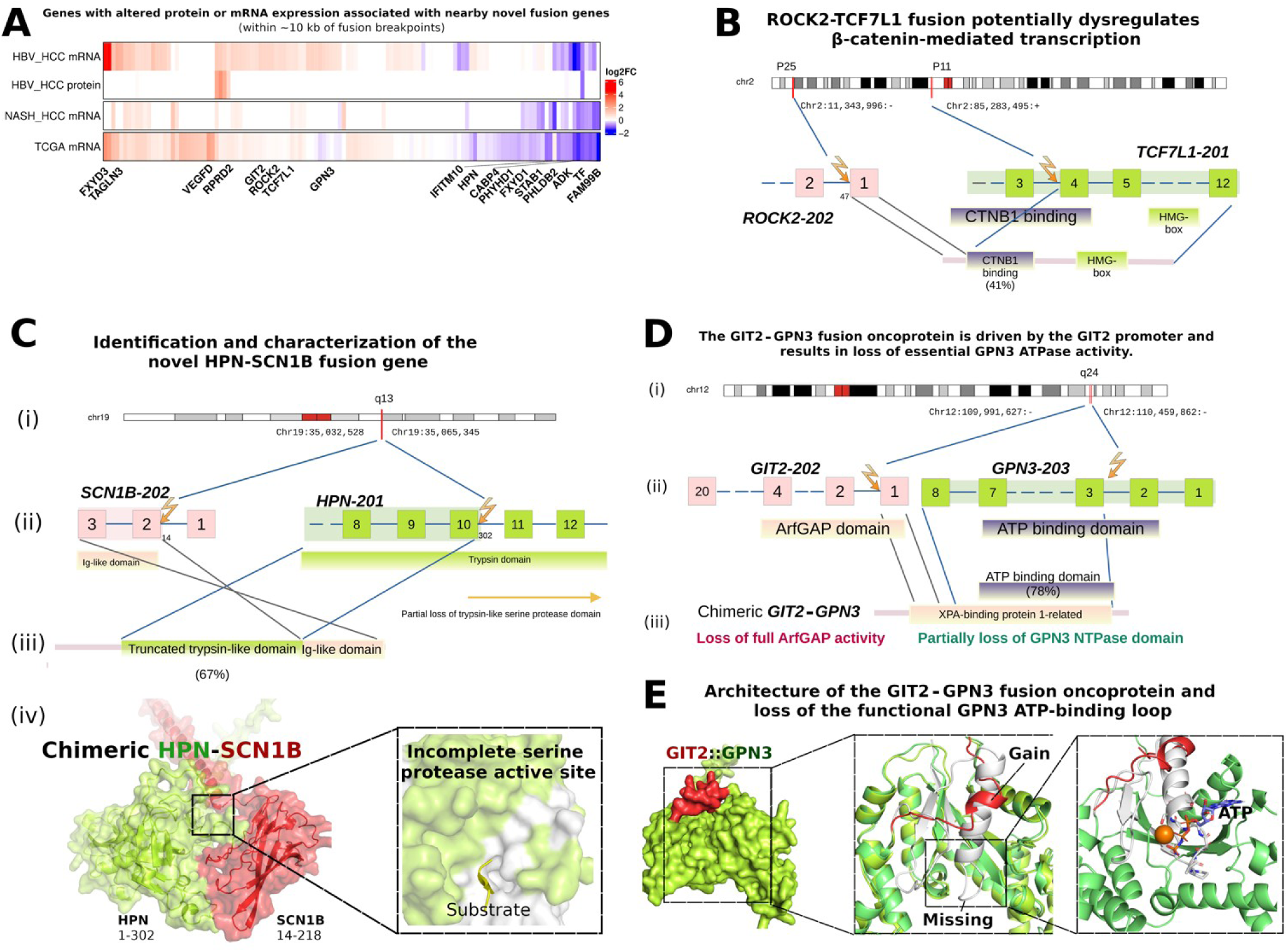
Fusion-associated gene expression and structural characterization of chimeric proteins in HCC. (**A**) Heatmap showing genes with significant expression differences between tumor and matched nontumor samples in mRNA (TCGA, MASLD-HCC, and HBV-HCC) or protein (HBV-HCC) datasets. Only genes located within ∼10 kb of novel tumor fusion breakpoints are included. Columns represent genes, with labels shown only for significantly altered genes (adj. P < 0.05 and log2FC > 2 or log2FC ← 1). Column clustering reveals recurrent patterns of fusion-associated expression alterations in hepatocellular carcinoma. (**B**) Schematic representation of the *ROCK2--TCF7L1* fusion generated by an intrachromosomal inversion on chromosome 2. Exon 1 of *ROCK2* is fused to exon 4 of *TCF7L1*, yielding an in-frame transcript. The resulting chimeric protein retains the N-terminal domain of ROCK2 (pink) linked to the truncated CTNNB1-binding domain and HMG DNA-binding domain of TCF7L1 (lemon green). (**C**) The HPN and SCN1B genes are both located on chromosome 19q13, and a genomic breakpoint fuses HPN exons 1-10 with SCN1B exons 2-3 to generate the novel HPN--SCN1B fusion gene. HPN normally comprises 13 exons, with the trypsin-like serine protease catalytic domain encoded by exons 8/9-13; the breakpoint truncates this domain but retains partial trypsin-like region of HPN. The SCN1B segment contributes its Ig-like domain, producing a chimeric protein that combines HPN-derived trypsin-like element with the SCN1B-derived Ig-like domain. Structural modeling illustrates the HPN regions in lemon green and the SCN1B regions in red, with the missing catalytic domain of HPN shown separately in complex with substrate (gold cartoon; PDB:1Z8G). (**D**) Schematic representation of the GIT2--GPN3 fusion generated by an intrachromosomal duplication on chromosome 12. The resulting in-frame transcript fuses the 5’ region of GIT2 to the 3’ region of GPN3. The chimeric protein retains a fraction N-terminal scaffolding region of GIT2 (pink), which is fused to a truncated P-loop NTPase domain of GPN3. (**E**) Structural overview of the chimeric GIT2--GPN3 fusion protein. The fusion is illustrated with the GIT2-derived segment in red and the GPN3-derived segment in lemon green. The schematic highlights the gain of the short, unstructured N-terminal peptide (residues 1-17) from GIT2 and the concomitant loss of the N-terminal region of GPN3 (residues 1-52). This truncation ablates a key loop (residues 11-18, highlighted) essential for coordinating ATP, thereby crippling the nucleotide-binding function of the GPN3 ATPase domain.

Together, these analyses position ADGRV1-TERT as an exemplary fusion oncogene that drives tumorigenesis while simultaneously yielding a stable, junction-specific candidate neoantigen for future evaluation in precision immunotherapy. At the same time, the SH3GLB2-PHYHD1 fusion demonstrates that fusion-derived neoantigens can be further diversified through PTMs.

### ICI-treated patients corroborate clinically-relevant fusion neoantigens

To identify patient-specific, clonally recurrent fusions and their associated neoantigens within a therapeutically relevant context, we analyzed the NCI-ICI cohort with patients undergoing active immunotherapy, defining 50 coding fusions (**Fig. 7A**). Notably, among ICI-treated patients the number of events was significantly higher in immunologically cold compared to hot tumors (Kruskal-Wallis test, p < 0.05; **Fig. S6a**). Subsequent immunopeptidomics revealed 33 potentially immunogenic peptides across fusion junctions (**Fig. 7B**). Still, to ensure therapeutic safety, we filtered peptides against healthy tissue proteomes, which identified KTHTGKPMK and LINLVNSSR with confirmed expression in normal human tissues (GTEx_32; **Table S3**). Consequently, 3 peptides were confirmed as endogenously presented only in clinical cancer proteomic datasets, providing direct evidence of their tumor-specific expression and establishing them as high-priority targets (**Fig. 7C, Table S3**).

**Figure 7.**
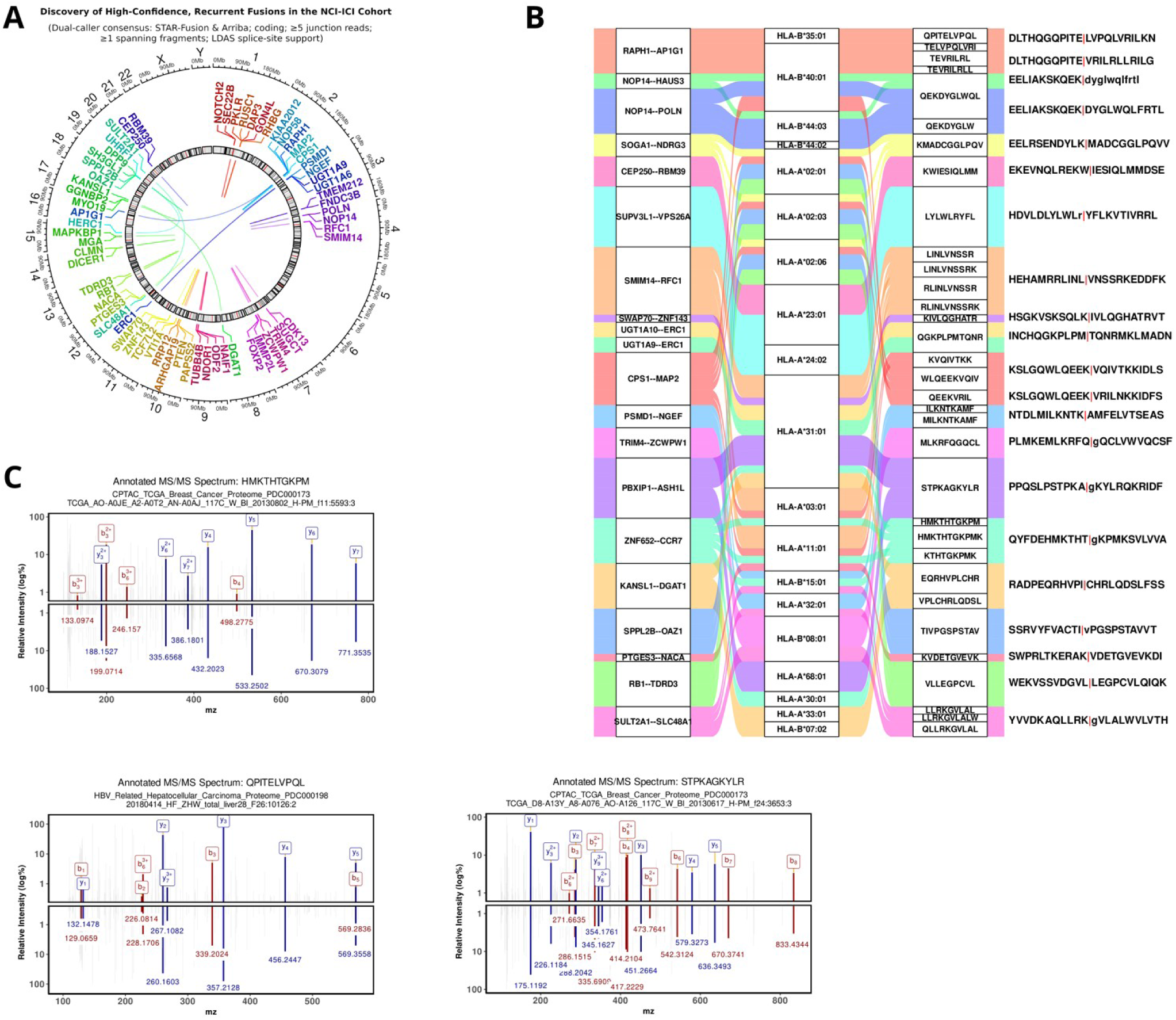
Fusion-derived immunogenic peptides identified in MASLD-HCC proteomes. (**A**) Circos plot of high-confidence gene fusions in the NCI-ICI cohort. Links connect chromosomal loci of coding, differentially expressed fusions identified in the NCI-ICIcohort. Only high-confidence fusions (≥5 junction reads, ≥1 spanning fragment, and splice-support criteria) are shown. (**B**) Alluvial diagram linking top fusion-derived immunogenic peptides to their originating fusion events, predicted MHC-I alleles, and 11-amino-acid flanking sequences. Fusion junctions are marked by a red vertical bar (‘|’); flow width reflects peptide detection frequency across samples. (**C**) Tandem mass spectra supporting endogenous expression of three fusion-derived candidate peptides detected in clinical cancer proteomes: HMKTHTGKPM (PDC000173), QPITELVPQL (PDC000198), and STPKAGKYLR (PDC000173).

## DISCUSSION

Although the present study establishes gene fusions as an abundant source of private neoantigens in MASLD-HCC, two conceptual controversies have emerged from recent advances in the field that challenge the direct translational applicability of our findings.

First, the assumption that MASLD-HCC constitutively benefits from neoantigen-driven immunotherapy stands in direct tension with mounting evidence of exceptional immunotherapy resistance in this specific etiology. Multiple large-scale reviews and meta-analyses now indicate that non-viral HCC, particularly MASLD/MASH-associated disease, exhibits significantly reduced sensitivity to immune checkpoint blockade compared to viral-associated HCC[39,40]. This diminished responsiveness has been mechanistically linked to a unique immunosuppressive tumor microenvironment characterized by lipid-laden Kupffer cells that promote T-cell exhaustion and dysfunction[40,41]. Moreover, recent single-cell atlas work has revealed that MASLD-positive HCC displays reduced tumor-reactive T-cell fractions despite increased total CD8⁺ T-cell infiltration, suggesting that the immune landscape is ‘cold’ and exhausted rather than primed for neoantigen recognition[42]. Our own finding that fusion burden correlates with an exhausted, low-cytolytic TME thus paradoxically places fusion-derived neoantigens, which are typically most effective in inflamed, T-cell-infiltrated tumors, into a context where their immunogenicity may be actively suppressed. This raises a critical translational question, if fusion-derived neoantigens in MASLD-HCC represent genuine candidate therapeutic targets, whose potency is undermined by the hostile TME or if the metabolic-immune interplay in MASLD fundamentally limits the utility of any neoantigen -based strategy. We therefore propose that fusion-derived neoantigen strategies are unlikely to achieve maximal efficacy as monotherapy and may instead require combination with therapies that restore T-cell function or remodel the tumor microenvironment, such as anti-TGF-β, anti-VEGF, or metabolic interventions. Mechanistically, the lipid-rich metabolic milieu of MASLD-HCC may impair effective recognition of fusion-derived neoepitopes by promoting T-cell exhaustion and dysfunction despite the presence of tumor-specific antigens[40,43,44], although this hypothesis requires experimental validation.

Second, despite proteogenomic integration, the field continues to face a persistent ‘precision gap’ between in silico prediction and biological reality of fusion-derived neoantigens. The latest advances underscore that DNA-level predictions alone remain inadequate, conventional pipelines relying on whole-exome sequencing and MHC binding algorithms achieve a positive predictive value of ≤5% for true immunogenic neoantigens, primarily because genomic variants do not reliably translate into presented peptides[45]. While our multi-tiered workflow including direct mass spectrometry confirmation represents a significant advance, recent data suggest that even mass spectrometry-validated presented peptides frequently fail to elicit functional T-cell responses, highlighting the additional layers of immune tolerance, T-cell receptor repertoire constraints, and tumor-intrinsic resistance mechanisms that separate immunopeptidomic identification from clinical efficacy. The ongoing phase I GT601 trial has attempted to address this by incorporating functional validation using autologous tumor-reactive T cells, achieving > 40% precision in identifying immunogenic neoantigens, yet remains restricted to the adjuvant setting after curative resection, a scenario that does not reflect the heterogeneous immune states typical of advanced MASLD-HCC[46]. Consequently, a central unresolved controversy is whether the field should prioritize ever-more-sensitive discovery pipelines to capture increasingly rare neoantigen classes, such as the phospho-epitopes described here, or whether limited clinical resources should instead be directed toward simplifying and accelerating validation for a narrower set of high-confidence antigens that can be rapidly deployed in combination with TME-modulating agents. The growing recognition that alternative splicing can produce ‘off-the-shelf’ neoantigens for HCC[47], potentially bypassing the need for fully personalized manufacturing, introduces yet another dimension to this debate. Accordingly, we envision a stratified translational framework in which recurrent fusion-derived candidate neoepitopes presented by common HLA alleles (e.g., ADGRV1--TERT in HLA-A*02:01-positive patients) may serve as “off-the-shelf” therapeutic candidates, whereas the majority of private fusion events will likely require personalized neoepitope selection based on patient-specific HLA haplotypes and fusion repertoires.

While our findings provide a compelling framework for targeting fusion events in MASLD-HCC, the path forward must reconcile the dual realities of a uniquely immunosuppressive TME and the persistent gap between neoantigen prediction and true immunogenicity. Addressing these controversies will require etiology-stratified clinical trials that prospectively compare neoantigen-based strategies across different HCC etiologies, alongside mechanistic studies that dissect how metabolic dysfunction directly impairs the presentation and recognition of fusion-derived epitopes. Furthermore, although the inclusion of the NCI-ICI cohort provides an opportunity to evaluate fusion-derived candidate neoepitopes in an immunotherapy-treated setting, the limited number of clinical responders precluded robust analyses of response or survival stratified by fusion status. Future studies in larger, prospectively annotated ICI-treated cohorts will be required to determine whether fusion-positive tumors are associated with improved clinical outcomes following immunotherapy. We present this work explicitly as a discovery pipeline and prioritized candidate resource, a necessary but not sufficient foundation for therapeutic success in this challenging disease.

## Supplementary Material

Supplementary text and figures.

## ACKNOWLEDGMENTS

The authors would like to thank the patients and their families for allowing research access to their samples. Also, we are grateful for access to all public data sets. The results are in part supported by data generated by the TCGA Research Network (https://www.cancer.gov/tcga) and European Genome Phenome Archive (EGA). L.N.Z would like to express her gratitude for support from Profs. Markus H. Heim, Hyungwon Choi, Matthew J. Watt, Philipp Kaldis, Qing Zhao (Lexie), and Patrik Midlöv.

## FUNDING

The laboratory of JBA is supported by competitive funding from the Novo Nordisk Foundation (0058419, 220C0074956, 0085704), Danish Cancer Society (R167-A10784, R278-A16638, R368-A21455), Independent Research Fund Denmark for Medical Research (4183-00118A, 1030-00070B).

## Author contributions

Conceptualization: L.N.Z and J.B.A; Data analysis and visualization: L.N.Z; Writing: L.N.Z and J.B.A. Funding: J.B.A.

## Data availability

The RNA-seq for healthy donor livers, MASLD and MASH patients have been deposited at GEO under accession number GSE281797. All clinical data, processed omics datasets, and relevant code related to MASLD cohort are accessible at https://github.com/SLINGhub/MASLD_dual_omics. The RNA-seq for MASLD-associated HCC has been deposited at GEO under accession number GSE334651.

## Competing interest

J.B.A declares consultancies for Flagship Pioneering, QED Therapeutics, and AstraZeneca. J.B.A has received funding from the Incyte Corporation and ADCendo but is not related to this study.

## Supplementary figure and table legends

Figure S1. Multi-modal characterization of recurrent fusion events in MASLD-HCC and pre-malignant liver tissue.

Figure S2. Comprehensive overview of high-confidence gene fusions in MASLD-HCC, including genomic locations, breakpoint patterns, differential expression, cross-cohort concordance, and snRNA-seq expression of fusion partner genes.

Figure S3. Transcriptional rewiring and immunogenic landscape of fusion events in MASLD-HCC.

Figure S4. The association between fusion event burden and tumor microenvironment remodeling in HCC.

Figure S5. Detection of endogenous peptides from MASLD-HCC associated gene fusions in clinical cancer proteomes.

Figure S6. Fusion-derived core epitopes and their MHC-I presentation in MASLD-HCC and public cancer proteomes.

Figure S7. Fusion-derived phosphopeptides as potential neoantigens and their clinical distribution.

Table S1: Top-tier recurrent fusion events across the entire cohort.

Table S2: Novel fusion events identified in the MASLD-HCC subset.

Table S3: Patient-specific, clonal fusions with immunogenic potential from the NCI-ICI cohort.

