## Supplementary figure for "Fusion-derived phospho-neoepitopes define a prioritized candidate neoantigen repertoire in MASLD-HCC"

### Neoantigen Landscape of Fusion Events Informs Tailored Immunotherapy for MASLD-HCC

**Li Na Zhao** and **Jesper B. Andersen**

*Supplementary materials*

#### Public datasets

We analyzed bulk RNA-seq data from 50 paired hepatocellular carcinoma samples[1] (GSE77314) and data from the National Translational Science Network of Precision-Based Immunotherapy for Primary Liver Cancer (phs003074.v1.p1; NCI-ICI) dataset[2], snRNA-seq data from MASLD-HCC tumors[3] (GSE189175), scRNA-seq datasets from two independent HCC cohorts[4,5] (GSE149614 and GSE189903).

This study incorporates a broad set of publicly available mass spectrometry–based proteomic datasets used for fusion detection, neoantigen discovery, and validation across multiple cancer types. These include CPTAC Proteomic Data Commons studies of HNSCC (PDC000221) [6],, ovarian cancer proteome (PDC000113, PDC000114) [7], ovarian cancer phosphoproteome (PDC000115) [8], pancreatic ductal adenocarcinoma (PDC000270) [9], breast cancer proteome (PDC000120) [10], gastric cancer proteomes (PDC000214, PDC000215) [11], clear cell renal cell carcinoma (PDC000127) [12], glioblastoma (PDC000204) [13], lung adenocarcinoma (PDC000153, PDC000224) [14,15], endometrial carcinoma (PDC000126) [16], colon cancer proteome (PDC000111) [17,18], and the CPTAC Prospective Breast BI Acetylome dataset (PDC000239) [10]. The CCLE proteome dataset (MSV000085836) supports additional neoantigen validation, while the NCI-ICI cohort immunopeptidomics datasets (PDC000173, PDC000198) [19,20] provide endogenous fusion-derived peptide evidence. The SH3GLB2–PHYHD1 fusion peptide analysis is supported by dataset PDC000232 [21]. Collectively, these datasets span proteome, phosphoproteome, acetylome, and immunopeptidome analyses across diverse cancers and cohorts.

#### RNA-seq of MASLD and MASLD-HCC cohorts (in-house data sets)

RNA-seq of MASLD cohort was performed by Biomarker Technologies (BMK) GmbH. Total RNA was extracted from liver biopsies using TRIzol reagent. RNA quantity and purity were assessed using a NanoDrop spectrophotometer, and RNA quality was evaluated using a Labchip GX system. cDNA libraries were prepared using the Hieff NGS Ultima Dual-mode mRNA Library Prep Kit for Illumina (Yeasen) according to the manufacturer's protocol. For MASLD-HCC, total RNA was isolated and shipped to Novogene. Both cohorts are carried out with paired-end RNA-sequencing (2×150 bp) on a Illumina NovaSeq 6000 platform. Raw sequencing data were quality-controlled with FastQC (v0.11.9) [22].

**Identification and filtering of high-confidence gene fusions**

RNA-seq data from in-house MASLD[23], MASLD-HCC cohort, and GSE77314[1], National Translational Science Network of Precision-Based Immunotherapy for Primary Liver Cancer (phs003074.v1.p1; NCI-ICI) dataset[2] were aligned to the GRCh38 reference genome using STAR (v2.7)[24]. Gene fusions were identified with two independent algorithms, STAR-Fusion v1.15.0[25] and Arriba v2.2.0[26], to increase robustness. Only high-confidence events were retained, defined as fusion fragments per million (FFPM) > 0.1 for STAR-Fusion and >1 split read support for Arriba.

To reduce false positives, multiple filtering steps were applied: (i) removal of fusions involving uncharacterized loci (e.g., lncRNAs annotated as ENSG IDs); (ii) exclusion of immunoglobulin (IGH/IGK/IGL) and mitochondrial (MT-) fusions; (iii) removal of isoforms of the same gene (geneA-genaA fusions); (iv) elimination of gene pairs previously detected in normal tissues, including TCGA normal samples, GTEx, or published non-cancer cell datasets; (v) Removal of artifacts: Fusions involving highly expressed liver-specific genes (e.g., *ALB*, *APOA2*, *HP*), pseudogenes, ribosomal or mitochondrial genes, or known read-through events were excluded.

Filtered candidate events were then cross-validated across both fusion callers and both cohorts. Only fusions reproducibly detected by STAR-Fusion and Arriba and present in multiple tumor samples were prioritized. Known fusion events and their oncogenic annotations were retrieved from FusionGDB2[27] to assess coding potential, functional class, and prior cancer relevance.

**IEDB-based MHC-I binding and processing predictions**

In complement to Trypsin Digestion and MHC Binding Assessment of Fusion Junctions, we also evaluate the immunogenic potential of peptides spanning fusion junctions identified from shared in-frame gene fusions detected by both Arriba and STAR-Fusion using the IEDB Analysis Resource (https://nextgen-tools.iedb.org/) to perform MHC-I binding and elution predictions, incorporating parameters such as class I pMHC immunogenicity, proteasome cleavage, TAP transport, and maximum precursor extension.

This approach allowed us to evaluate the processing likelihood of candidate peptides and further refine the list of high-potential immunogenic epitopes.

**Cross-reactivity and immunogenicity prediction**

To refine the list of candidates that present fusion-derived neoantigens, only peptides detected in cancer proteomes were retained, excluding those identified in normal datasets. Potential cross-reactivity was assessed by aligning each 8-11 amino acid peptide against the human reference proteome using the Ensembl BLASTP tool[28] to identify identical or near-identical self-peptides; no significant matches were found for epitopes detected in clinical cancer proteomes. To evaluate potential pre-existing immune recognition, all peptides were queried against experimentally validated epitopes in the IEDB (https://www.iedb.org/) and TCRmatch databases[29]. Expression profiles of source genes corresponding to any matched self-peptides were further examined across normal tissues using GTEx[30] and the Human Protein Atlas[31] to assess off-target risks.

**Neoepitope pMHC-TCR complex modeling**

To model the YTMRLWTQAR neoepitope within its HLA context, we systematically surveyed 1,036 experimentally solved 3D immune complexes (including antibody-antigen, TCR-pMHC, and MHC-ligand structures) available in the Immune Epitope Database (IEDB) and Protein Data Bank (PDB) as of October 30, 2025, encompassing 672 unique core epitopes across diverse HLA class I alleles. The predicted HLA restrictions for YTMRLWTQAR, HLA-A68:02, HLA-A68:01, HLA-A31:01, HLA-A33:01, and HLA-A11:01, were used to filter potential structural templates. Sequence similarity between YTMRLWTQAR and known MHC-bound epitopes was quantified using pairwise alignment based on the BLOSUM62 substitution matrix, with gap opening and extension penalties applied. Structural templates were prioritized according to (i) HLA allele compatibility, with emphasis on HLA-A68:01 and HLA-A11:01, and (ii) the highest BLOSUM62 similarity scores*[32]* reflecting physicochemical resemblance to the query peptide. This analysis identified ETSPLTAEKL-HLA-A68:01 (PDB: 6EI2) and SALEWIKNK-HLA-A11:01 (PDB: 7S8R) as the most relevant templates for modeling the YTMRLWTQAR-pMHC complex. The TCR-pMHC complex with PDB code 8RYQ (Structure of S8-9F3 TCR in complex with HLA-A11:01 bound to the ELFSYLIEK peptide) was selected as the primary template to model the TCR docking geometry and orientation relative to the predicted YTMRLWTQAR-HLA-A*11:01 complex. All homology modeling and complex reconstruction steps were performed using MODELLER.

**Stringent filtering criteria**:

- Removed: Extremely high-frequency fusions (TVP23C--CDRT4, A1BG-AS1--APOA2)
- Excluded liver-specific expression artifacts: Albumin (ALB), apolipoproteins (APO), olfactory receptors (OR), and fibrinogen (FGB/FGG)
- Filtered common/classic technical chimeras: Ribosomal proteins (RP), heat shock proteins (HSP serum proteins (SAA, SERPIN)
- Removed metabolic enzymes: Liver-abundant genes (CPS1, FTCD, PKLR) without clear cancer relevance
- Antisense or pseudogenes (MPHOSPH10P3, PRSS42P, etc.)
- Excluded non-coding RNAs: LINC* and MIR*HG pseudogenes without mechanistic rationale

#### The unfiltered fusion landscape

Initial Recurrence Analysis Identifies both Putative Drivers and Prevalent Artifacts: Our initial, unfiltered analysis of recurrent gene fusions revealed a heterogeneous landscape across liver cancer cohorts (Fig. S1a). Within MASLD-related cohorts[33], we observed a significant number of recurrent fusions, suggesting a conserved pattern of genomic rearrangements (Fig. S1b). Strikingly, the most frequently detected events were dominated by fusions involving genes with high, liver-specific expression, such as ALB (albumin), APOC2, and SAA1 (Fig. S1c).

Metabolic and Liver-Specific Fusions are Detectable in Pre-Cancerous Tissue: Several of these highly recurrent fusions were detectable not only in tumors but also in histologically non-tumor liver tissue from MASLD patients (Fig. S1b), indicating they can arise early in disease progression. For instance, fusions involving the ALB locus were present in 53% (46/87) of pre-cancerous MASLD samples. Their presence was not associated with any of the 63 clinical, laboratory, or histological variables assessed, nor with a reduction in serum albumin levels (Fig. S1d), suggesting they are not simple markers of liver dysfunction.

Transcriptomic Evidence Supports an Artifact Model for Prevalent Fusions: In contrast to high-confidence, DNA-level fusions which often show nuclear retention of the primary transcript, these recurrent fusions exhibited balanced expression of both partner genes in snRNA-seq data (GSE189175) [3,34,35]. Furthermore, fusions like APOC2--SAA1, ALB--FGB, and SERPINA1--ALB were consistently detectable in both nuclear (snRNA-seq; GSE189175) and cytoplasmic (scRNA-seq; GSE149614 [4,36], GSE189903 [37]) RNA pools (Fig. S1e). This expression pattern is more consistent with trans-splicing or read-through transcription between co-expressed genes, rather than a DNA-level rearrangement.

Rationale for Subsequent High-Confidence Filtering: The prevalence of these likely expression-associated artifacts motivated the implementation of a stringent, dual-caller filtering pipeline to distinguish them from genuine, DNA-level oncogenic drivers.

#### Trypsin digestion and MHC binding assessment of fusion junctions

From the identified set of shared in-frame gene fusions that were detected by both Arriba and STAR-Fusion. We extracted the peptide sequences spanning the fusion junctions. Using the cleave function [38], we simulated trypsin digestion of these junction peptides to generate all possible cleavage fragments. The resulting peptides were then evaluated for potential immunogenicity using MHCpan I (netMHCpan-4.2) [39] and II (NetMHCIIpan-4.3) [40], identifying strong binders across a representative set of five common human MHC alleles (HLA-A*01:01, HLA-A*02:01, HLA-A*24:02, HLA-B*07:02, and HLA-B*51:01). This analysis produced a filtered list of fusion-derived peptides containing potential MHC class I and II binding epitopes, which serve as candidates for further immunogenicity assessment and CAR-T target development.

#### Trypsin-based fusion neoepitope screen

First, trypsin digestion followed by MHC binding assessment identified 13 fusion events (9 high-confidence) whose junction-derived peptides are predicted to strongly bind MHC-I (SB; ≤0.5%; Fig. S4a; Table S1), along with 40 additional peptides predicted as weak binders (WB; 0.5-2%; Fig. Sbc; Table S1). For these 16 fusion-derived strong binders exhibit high predicted MHC-I binding (EL percentile < 2), positive immunogenicity scores, efficicent proteasomal processing and adequate TAP transport (NetCTLpan combined score > 0.5). Furthermore, 5/16 are experimentally detected across multiple cancer types (Fig. S4c, Table S1): KLRGPDVFSSK in ovarian cancer (PDC000113/114); LRILLQSKNVM in breast cancer (PDC000173); SPRKILTM and GPQVKSRLAF (acetylated at K5) in HBV-related hepatocellular carcinoma (PDC000198); and YLLYCPCMV in colon cancer (PDC000111) [17,18]. Notably, the detection of GPQVKSRLAF with lysine-5 acetylation represents a post-translationally modified neoantigen candidate. These findings demonstrate that fusion-derived peptides are endogenously processed and presented in vivo, highlighting their potential as immunogenic neoantigen candidates for cancer immunotherapy.

##### Fusion transcript abundance

Fusion transcript abundance was estimated by calculating the fusion fraction, defined as the ratio of fusion transcript expression to the expression of the corresponding partner transcripts. Fusion expression was quantified as fusion fragments per million (FFPM) using STAR-Fusion and normalized to isoform-level transcript abundance (TPM) generated by RSEM. Fusion fractions were calculated using two complementary approaches: (i) relative to the combined expression of both partner genes and (ii) a promoter-aware approach based on the transcriptionally active partner gene. These metrics were used to characterize the relative abundance of fusion transcripts and were not used to infer DNA-level clonality.

##### Estimation of fusion clonality and tumor mutational burden

Matched tumor-normal whole-exome sequencing (WES) data (NCI-CLARITY study; phs003074) were analyzed using GATK Mutect2 (v4.6.1.0)[41]. High-confidence somatic variants were retained by selecting PASS variants and annotated with Funcotator. Variant allele frequencies (VAFs) were extracted from the filtered somatic VCFs, and coding nonsynonymous variants were classified using an operational VAF threshold, where variants with VAF $\geq 0.30$ were considered clonal and variants with VAF $<0.30$ were considered subclonal. Tumor mutational burden (TMB) was calculated from filtered coding nonsynonymous variants using the pyTMB[42] pipeline and normalized to an effective exome size of 33.28 Mb. Fusion transcripts were independently identified from RNA-seq using STAR-Fusion and quantified as fusion fragments per million (FFPM). RNA-seq was used to detect and quantify fusion transcripts, whereas clonal and subclonal classifications were derived from matched WES data.

##### Tiered immunoinformatic filtering reveals a high-confidence repertoire

To identify therapeutically relevant fusion-derived neoantigens, we implemented a multi-tiered computational pipeline. First, we extracted in-frame fusion junctions and generated an 8-amino-acid window spanning the breakpoint for each event. Concurrently, we performed a comprehensive MHC-I immunogenicity prediction on all possible 9-10mer peptides, retaining only high-confidence candidates with a NetCTLpan combined score $> 0.8$ (indicating efficient antigen processing) and an MHCflurry percentile rank $\leq 2.0$ (indicating strong MHC binding affinity). These high-confidence binders were then cross-referenced against our library of fusion junction sequences to pinpoint epitopes spanning the novel breakpoint. This integrated bioinformatic workflow successfully identified 158 unique neoepitopes derived from 30 distinct fusion events, providing a prioritized list of candidates for downstream functional validation.

#### Patient-matched fusion neoantigens

Our integrated computational and proteomic pipeline successfully identified patient-specific, HLA-matched fusion neoantigens in the MASLD-HCC cohort [43]. We detected four distinct fusion events across three patients, each yielding high-confidence neoepitopes peptides with their cognate HLA alleles, supported by RNA-level expression evidence. Patient R97 presented two independent fusion-derived targets (*ALDH2--ACAD10* and *CUX2--CABP1*), each generating a single immunogenic epitope (GPQVKSRLAF and DLRRLQDRSL, respectively) with moderate to high expression levels (FFPM 0.34-0.62). Patient R59 harbored a *BRD9--ABCA3* fusion producing two distinct but overlapping epitopes (SMLSSLGYI; ISMLSSLGY), while patient R26 exhibited a highly expressed *SRSF10--MRPS21* fusion (FFPM 1.69) generating one immunogenic peptide (HSDNDRILTM). These findings demonstrate the feasibility of identifying therapeutically relevant, patient-specific fusion neoantigens in MASLD-HCC, with several patients presenting multiple targetable events.

#### YTMRLWTQAR detection

YTMRLWTQAR was detected in both its phosphorylated (pT7) and unphosphorylated forms within tumor samples. The MS/MS spectrum of the phosphorylated peptide exhibited suppressed fragmentation C-terminal to T7, a well-documented phenomenon for phosphopeptides where the labile phosphate group dominates the fragmentation energy [44,45]. Despite this, we confidently localized the modification to T7 through detected flanking b- and y-ions. The presence of both forms suggests that phosphorylation at T7, a residue predicted to be a key TCR contact, may act as a dynamic regulatory switch. This is supported by studies showing that phosphorylation can critically alter peptide conformation in the MHC groove and that TCRs can be exquisitely specific for the phosphorylated form, fundamentally changing immunogenicity [46,47]. This dynamic modification could thus represent a mechanism for immune evasion or antigenic diversification [47].

#### ADGRV1-TERT fusion junction identification

The ADGRV1-TERT fusion peptide (LLIVILKGIYHQSMSQIYGLIHGDLSYVQCQGIPQGSILSTLLCSLCYGD) was confidently detected across multiple CPTAC proteomic datasets, including clear cell renal cell carcinoma (CCRCC, PDC000127), glioblastoma (GBM, PDC000204), lung adenocarcinoma (LUAD, PDC000153 and PDC000224), uterine corpus endometrial carcinoma (UCEC, PDC000126), and gastric cancer

(PDC000215) phosphoproteome studies. Peptide-spectrum matches (PSMs) were consistently assigned a "Yes" confidence status, with observed post-translational modifications including oxidation of methionine (M@14), methylation of lysine (K@7), and carbamidomethylation of cysteine residues (C@30, C@44, C@47). The peptide was detected across both proteome and phosphoproteome datasets, demonstrating reproducible identification in diverse tumor contexts and analytical workflows. These findings support the robustness and biological relevance of this TERT fusion-derived peptide (PDC000126, PDC000127, PDC000153, PDC000204, PDC000215, PDC000224).

#### Neoantigen prediction and validation

Neoantigens derived from the ADGRV1-TERT fusion junction were predicted using the Next-Generation IEDB Analysis Resource tools. Putative epitopes were filtered for strong MHC-I binding (IC50 < 500 nM, percentile rank ≤ 2) and a positive immunogenicity score. The physiological presentation of the top candidate was validated by searching for the peptide sequence in mass spectrometry data from the CPTAC Prospective Breast Cancer Acetylome (PDC000239) and other public CPTAC datasets. A peptide-spectrum match (PSM) p-value < 0.05 was used as the criterion for confident identification.

#### Supporting Figures and Text

#### Figure S1

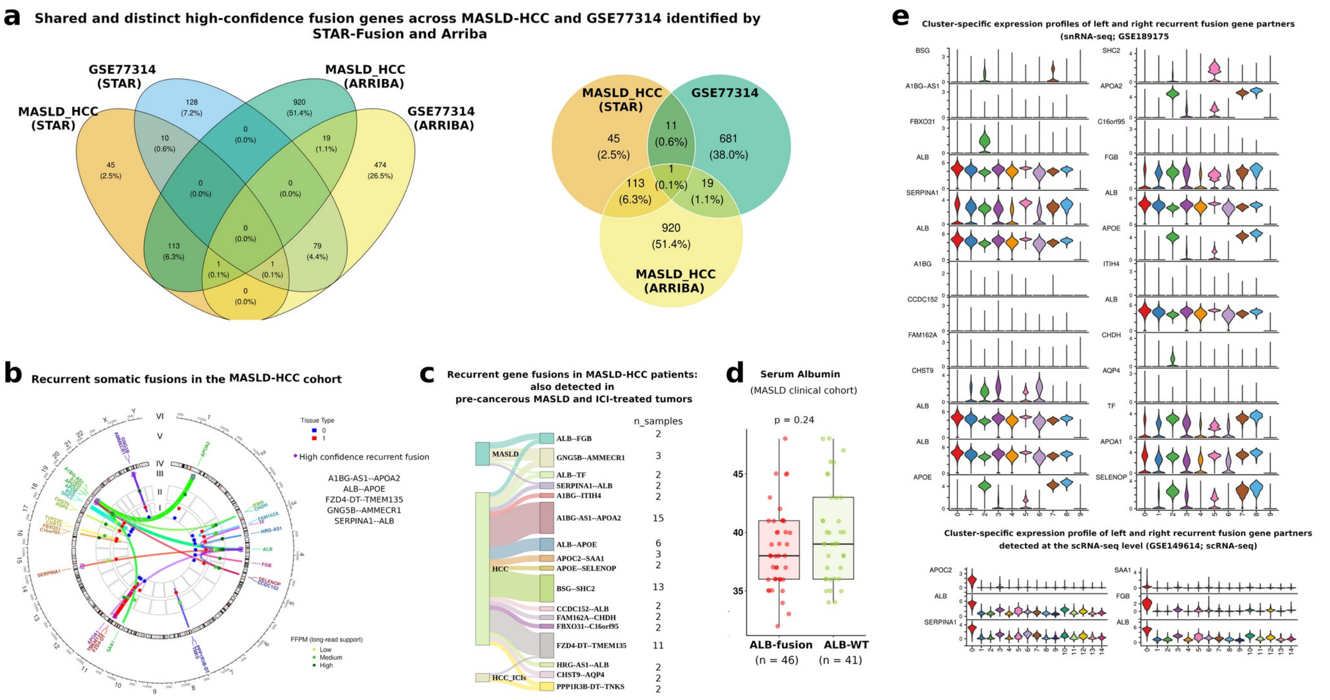

**Figure S1. Multi-modal characterization of recurrent fusion events in MASLD-HCC and pre-malignant liver tissue.**

(a) Left: Venn diagram illustrating the overlap of filtered fusion events between the MAFLD-HCC dataset and the GSE77314 cohort, as detected by STAR-Fusion and Arriba. Fusion events were filtered to exclude uncharacterized or non-coding genes (identified by ENSG-based names), immunoglobulin-related fusions (IGH, IGK, IGL), mitochondrial gene fusions (MT-), and same-gene fusions (e.g., GENE--GENE). Only high-confidence events with FFPM > 0.1 were retained. Right: Venn diagram illustrating the overlap of fusion genes identified in MAFLD-HCC samples using two detection approaches and a public liver cancer dataset (GSE77314). Notably, the fusion MTAP--CDKN2B-AS1 is consistently detected in both MAFLD-HCC and GSE77314, suggesting it may represent a recurrent or biologically relevant event in liver cancer. Differences across sets highlight method-specific and disease-context fusion profiles.

(b) Circos plot showing the recurrent fusion landscape in the MAFLD-HCC cohort. Starting from the inner circle to the outer: connections of somatic gene fusion partners; (I) tissue type, colored red for tumor and blue for non-tumor; (II) coverage of supported reads by long-read transcriptome sequences (FFPM), colored from warm to cold (yellow green to dark green) to reflect increasing read support; (III) high-confidence fusions, shown as purple diamonds; (IV) chromosome ideogram providing genomic context; (V) gene label of fusion pairs, and (VI) chromosome labels. This arrangement highlights the recurrence, confidence, and tissue specificity of somatic fusions.

(c) Recurrent gene fusion events identified in MASLD-HCC tumors and their presence in pre-cancerous MASLD tissue. Line width corresponds to recurrence frequency (number of samples). The ALB locus emerges as a genomic hotspot with multiple fusion partners. GNG5B::AMMECR represents the most frequently shared fusion between pre-malignant and malignant stages, suggesting its role as an early driver event in hepatocarcinogenesis.

(d) Comparison of serum albumin level in pre-cancerous MASLD cohorts stratified by ALB fusion status. Boxplots show median and interquartile ranges, with individual data points overlaid. Statistical significance was assessed using the Wilcoxon rank-sum test (p-values shown).

(e) Stacked violin plots depicting the expression distributions of the Left Gene (left) and Right Gene(right) of recurrent fusion partners across cell clusters identified by Seurat (Top: snRNA-seq; GSE189175; Bottom: scRNA-seq; GSE149614). Expression values are grouped by cluster identity, with violin width reflecting cell density at a given expression level.

#### Figure S2

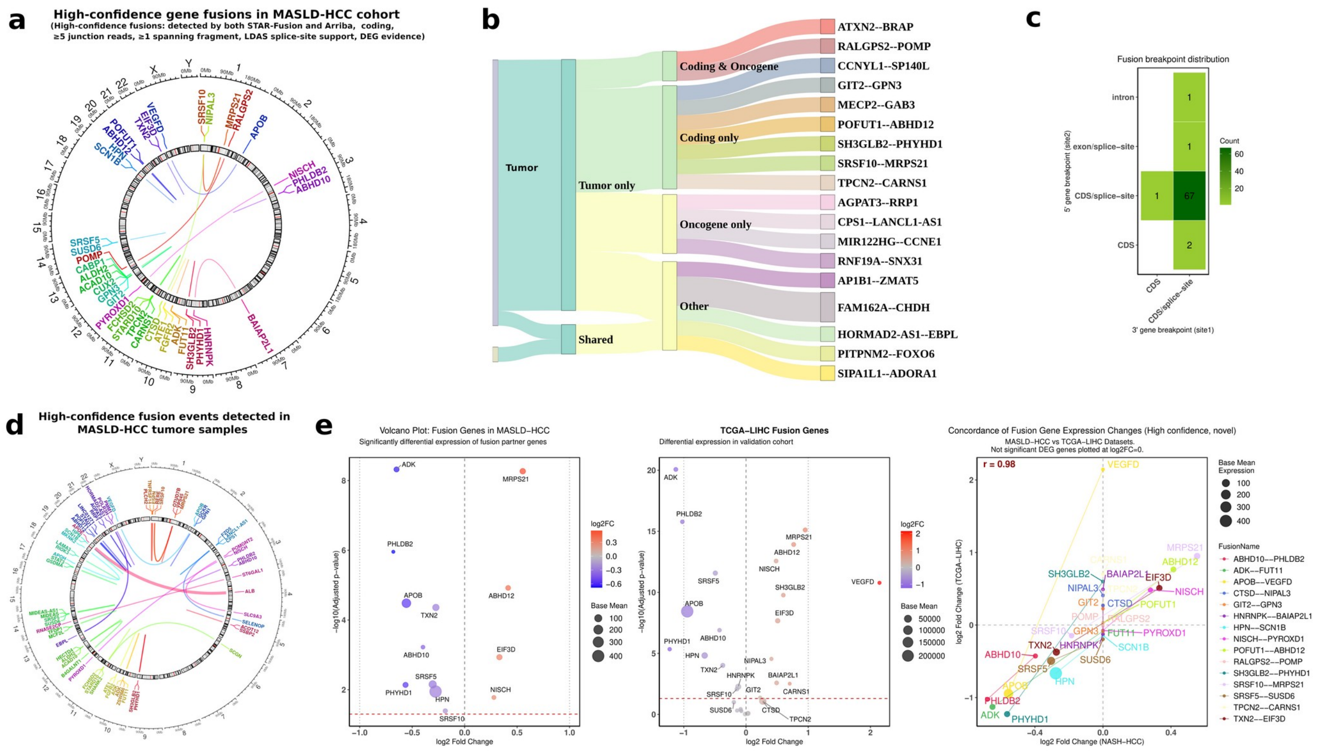


**Figure S2. Comprehensive overview of high-confidence gene fusions in MASLD-HCC, including genomic locations, breakpoint patterns, differential expression, cross-cohort concordance, and snRNA-seq expression of fusion partner genes.**

(a) Circos plot of high-confidence gene fusions in the MASLD-HCC cohort (including both novel and known). Only fusions detected by both STAR-Fusion and Arriba in tumor samples, with ≥5 junction reads, ≥1 spanning fragment, and supported by canonical or inclusive non-reference splice sites, are shown. Chromosomes (hg38) are displayed as an outer ideogram, with gene labels and links colored by fusion pair. Curved links connect fusion partners, with width scaled by the number of supporting samples. The plot highlights high-confidence fusion events, multi-transcript support, and both intra- and inter-chromosomal rearrangements.

(b) This Sankey network visualizes the distribution of high-confidence fusion gene events detected by both STAR-Fusion and Arriba across liver tumor and non-tumor tissues. Fusion calls were filtered based on expression (FFPM > 0.1), junction read support (≥ 5), anchor support, and known oncogene or database annotations to ensure reliability.

(c) Heatmap of fusion breakpoint locations showing that most in-frame fusions occur at CDS or CDS/splice-site regions in both partner genes.

(d) Circos visualization of high-confidence gene fusions in MASLD-HCC tumor samples. The plot shows coding, differentially expressed (DEG-supported) gene fusions. Only high-confidence fusions (≥5 junction reads, ≥1 spanning fragment, and splice-support criteria) with evidence of high junction read counts are shown.

(e) Left: Differential expression of fusion partner genes in MASLD-HCC. Scatter plot shows $\log_2$ fold change (x-axis) versus $\log_{10}$(base mean expression) (y-axis) for genes involved in high confidence, novel fusion events. Genes with positive $\log_2$ fold change are upregulated in tumor samples relative to non-tumor controls. The plot highlights which fusion partners are significantly dysregulated in the MASLD-HCC cohort. Middle: Significantly differential expression of fusion partner genes in TCGA-LIHC. Right: Concordance of fusion gene expression changes between MASLD-HCC and TCGA-LIHC cohorts.

**Figure S3**

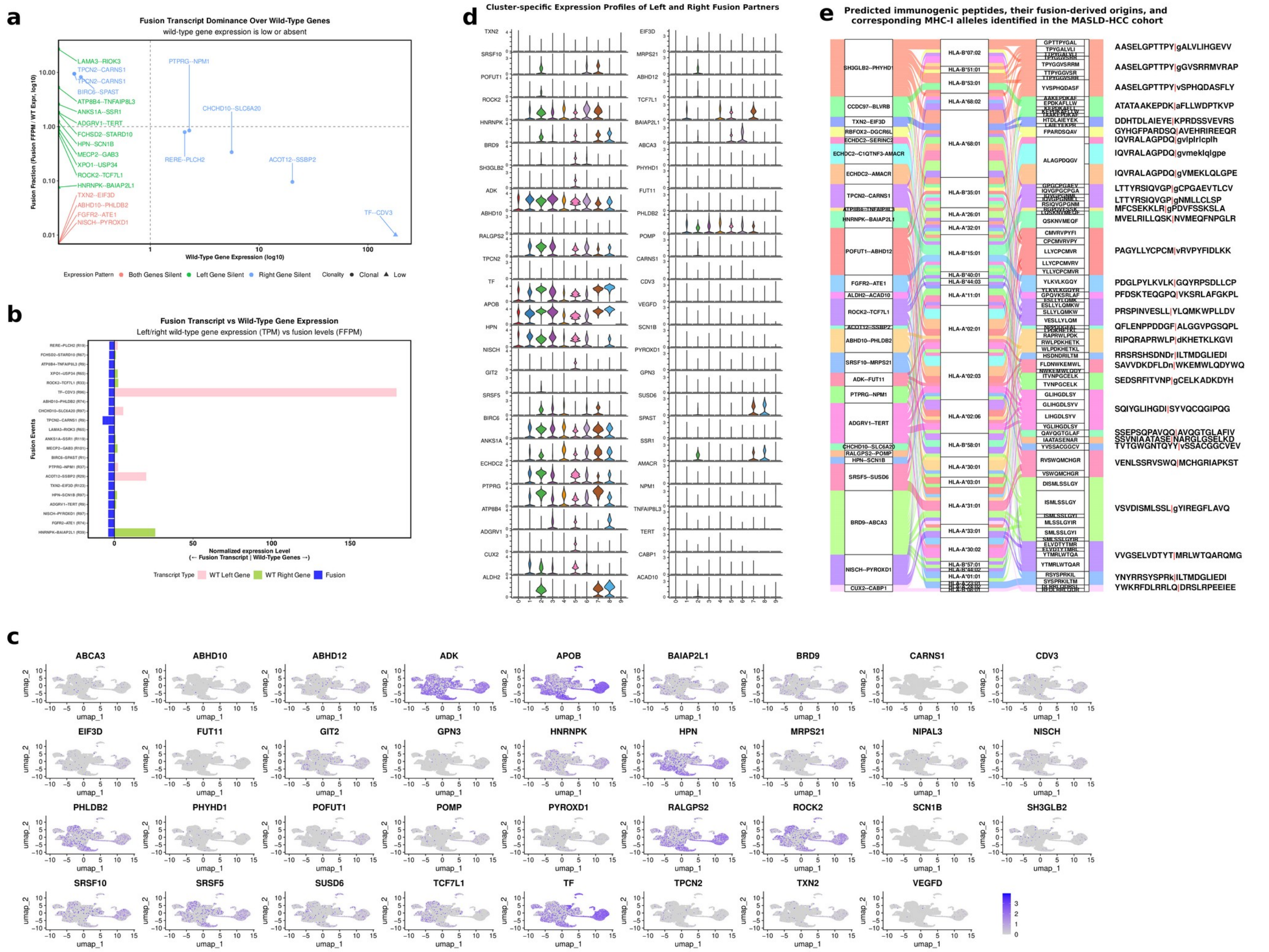

**Figure S3. Transcriptional rewiring and immunogenic landscape of fusion events in MASLD-HCC.**

(a) Scatter plot showing fusion fraction relative to wild-type expression, colored by silencing pattern and shaped by clonality.

(b) Diverging bar plot comparing absolute expression levels of 22 fusion transcripts versus wild-type partner genes. Both (a) and (b) highlight the transcriptional rewiring where fusion events become the dominant species, particularly in contexts of wild-type gene silencing.

(c) Feature plots showing expression of genes involved in high-confidence novel fusion events detected by snRNA-seq.

(d) Stacked violin plots depicting the expression distributions of the Left Gene (left) and Right Gene(right) of fusion partners across cell clusters identified by Seurat (snRNA-seq; GSE189175). Expression values are grouped by cluster identity, with violin width reflecting cell density at a given expression level.

(e) Alluvial representation of top fusion-derived immunogenic peptides in the MASLD-HCC cohort. Each flow connects fusion events to their corresponding peptides, predicted MHC-I alleles (allele), and the 11-amino-acid left and right sequences flanking the fusion junction. Peptide junctions are highlighted with a bold red vertical bar (‘|’). The width of each flow reflects the frequency of peptide occurrence across patients or samples. Colors indicate different fusion events.

#### Figure S4

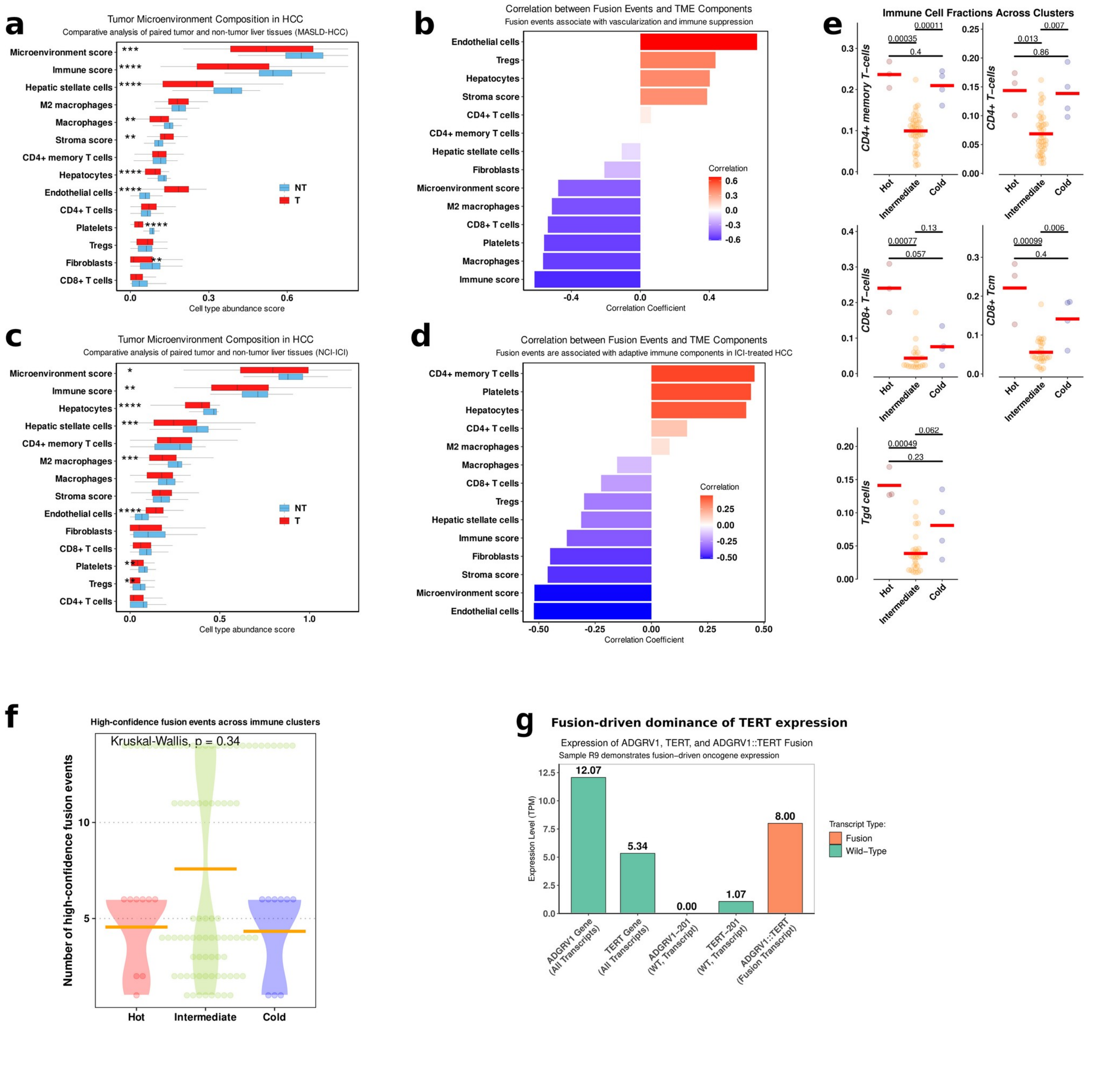


**Figure S4. The association between fusion event burden and tumor microenvironment remodeling in HCC.**

(a) Comparative analysis of tumor microenvironment composition in MASLD-HCC. Box plots display abundance scores for immune cells, stromal components, and composite scores derived from transcriptomic deconvolution of paired tumor (T) and non-tumor (NT) liver tissues from MASLD-HCC cohort. Cell types are ordered by median expression across all samples. Statistical comparisons were performed using Wilcoxon signed-rank test (*p < 0.05, **p < 0.01, ***p < 0.001).

(b) High confidence fusion events correlate with specific tumor microenvironment remodeling in MASLD-HCC**.** Spearman correlation analysis reveals fusion burden positively associates with endothelial cell abundance (vascularization) while negatively correlating with multiple immune components. Bars represent correlation coefficients (ρ) between fusion event counts and cell type abundance scores derived from transcriptomic deconvolution with xCell.

(c) Comparative analysis of tumor microenvironment composition in NCI-ICI.

(d) High confidence fusion events correlate with specific tumor microenvironment remodeling in NCI-ICI. Unlike the broad immune depletion observed in treatment-naïve HCC, ICI-treated tumors display selective alterations: fusion events correlate with increased $CD4^+$ memory T-cells ($\rho = +0.46$, $p = 1.2\times10^{-5}$) and platelets ($\rho = +0.44$, $p = 2.5\times10^{-5}$), while maintaining negative correlations with cytotoxic $CD8^+$ T-cells ($\rho = -0.22$, $p = 0.04$) and Tregs ($\rho = -0.30$, $p = 0.006$). This suggests that immunotherapy may partially rescue certain adaptive immune components while failing to reverse fusion-driven $CD8^+$ T-cell exclusion. Fusion events associate with reduced stromal scores ($\rho = -0.46$, $p = 1.0\times10^{-5}$) and fibroblast content ($\rho = -0.45$, $p = 1.7\times10^{-5}$), contrasting with the complex stromal alterations seen in treatment-naïve HCC.

(e) Distinct immune composition characterizes cold, intermediate, and hot MASLD-HCC microenvironments. Cellular enrichment scores derived from xCell deconvolution demonstrate significant differences in immune cell abundance across the three immune clusters defined by the Ayres 18-gene signature. The visualization combines individual sample points (beeswarm) with cluster means (red crossbars), highlighting both the distribution and central tendency of immune cell fractions. Statistical testing confirms significant differences in specific lymphocyte populations between clusters, consistent with the continuum of immune activation states in hepatocellular carcinoma.

(f) Fusion event burden is comparable across immune clusters in MASLD-HCC. Violin and beeswarm plots show the distribution of high-confidence fusion events across immunologically hot (red), intermediate (yellowgreen), and cold (blue) tumor clusters. Each point represents an individual tumor sample, with orange crossbars indicating mean fusion events per cluster. Statistical testing (Kruskal-

Wallis) revealed no significant differences in fusion burden between immune clusters, suggesting that total fusion quantity alone does not determine the immune contexture of MASLD-HCC tumors.

(g) Expression levels of ADGRV1, TERT, and the ADGRV1–TERT fusion in sample R9. Wild-type ADGRV1 and TERT transcripts are shown alongside the fusion transcript. Expression is measured in TPM (transcripts per million). The ADGRV1–TERT fusion is highly expressed, while wild-type TERT is largely silenced, highlighting the fusion as the dominant oncogenic transcript. Values above bars indicate exact TPM levels. Color distinguishes transcript type: wild-type (green) and fusion (orange).

**Figure S5**

### Detection of Endogenous Peptides from MASLD-HCC Associated Gene Fusions in Clinical Cancer Proteomes

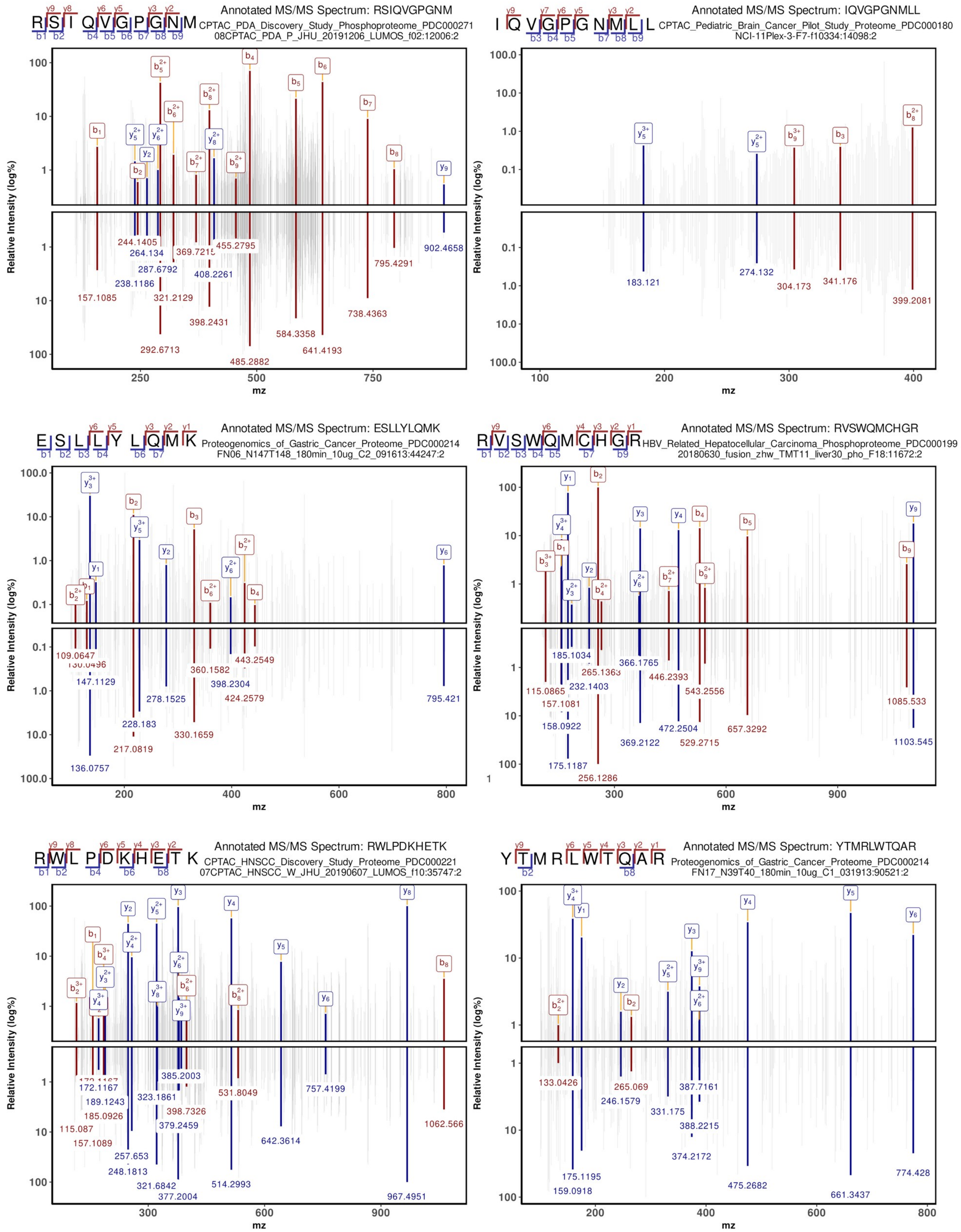

**Figure S5. Detection of endogenous peptides from MASLD-HCC associated gene fusions in clinical cancer proteomes.**

Annotated MS/MS spectra show endogenous peptides from MASLD-HCC gene fusions, with matched b-ions (blue) and y-ions (red) labeled by position and charge: RWLPDKHETK (ABHD10--PHLDB2) from HNSCC proteome (PDC000221); YTMRLWTQAR (NISCH--PYROXD1) and ESLLYLQMK (ROCK2--TCF7L1), both containing methionine oxidation, from gastric cancer proteome (PDC000214); RVSWQMCHGR (SRSF5--SUSD6) with carbamidomethylation from HCC phosphoproteome (PDC000199); and IQVGPGNMLL (TPCN2--CARNS1) and RSIQVGPGNM (TPCN2--CARNS1) from pediatric brain cancer (PDC000180) and PDA phosphoproteome (PDC000271), respectively. All spectra represent high-confidence identifications (PepQuery-validated 'Yes', PSM scores >12, mass tolerance = 0.06 Da). Note that SYSPRKILTM and YGLIHGDLSY, while receiving automated 'Yes' validations, were excluded due to manual curation revealing sparse fragmentation patterns with low peak counts and intensity, insufficient for definitive neoantigen confirmation.

**Figure S6**

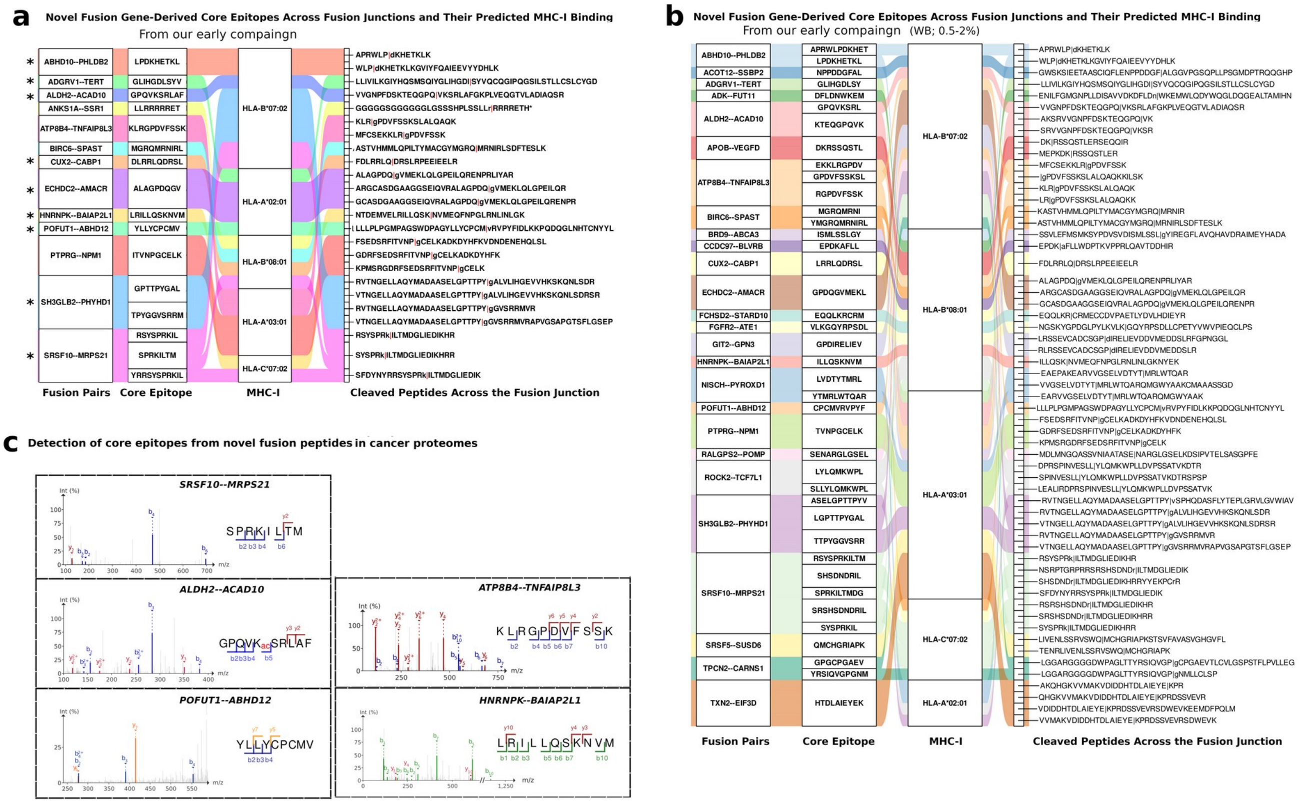

**Figure S6. Fusion-derived core epitopes and their MHC-I presentation in MASLD-HCC and public cancer proteomes.**

(a) Alluvial plot depicting the flow from novel fusion genes to core epitopes spanning the fused junction, their matched MHC-I presentation (strong binder), and cleaved peptide products in our early campaign. The notation is as follows: '|' marks the fusion breakpoint; lowercase letters indicate non-template bases (extra bases not in the reference), or, when within the coding sequence, reference mismatches (SNPs/SNVs); and '*' denotes a stop codon.

(b) Alluvial plot depicting the flow from novel fusion genes to core epitopes spanning the fused junction, their matched MHC-I presentation (weak binder), and cleaved peptide products in our early campaign.

(c) Detection of core epitopes from novel fusion peptides in cancer proteomes. Core epitopes (KLRGPDVFSSK, LRILLQSKNVM, SPRKILTM, YLLYCPCMV, GPQVKSRLAF) were onfidently identified across multiple public proteomics datasets (CPTAC TCGA Breast PDC000173, Colon PDC000111, Ovarian PDC000113_PDC000114, and HBV-related Hepatocellular Carcinoma PDC000198). MS/MS spectra of these peptides were inspected using PDV with a mass tolerance of 0.6 Da. Observed post-translational modifications include acetylation at K5 for GPQVKSRLAF.

**Figure S7**

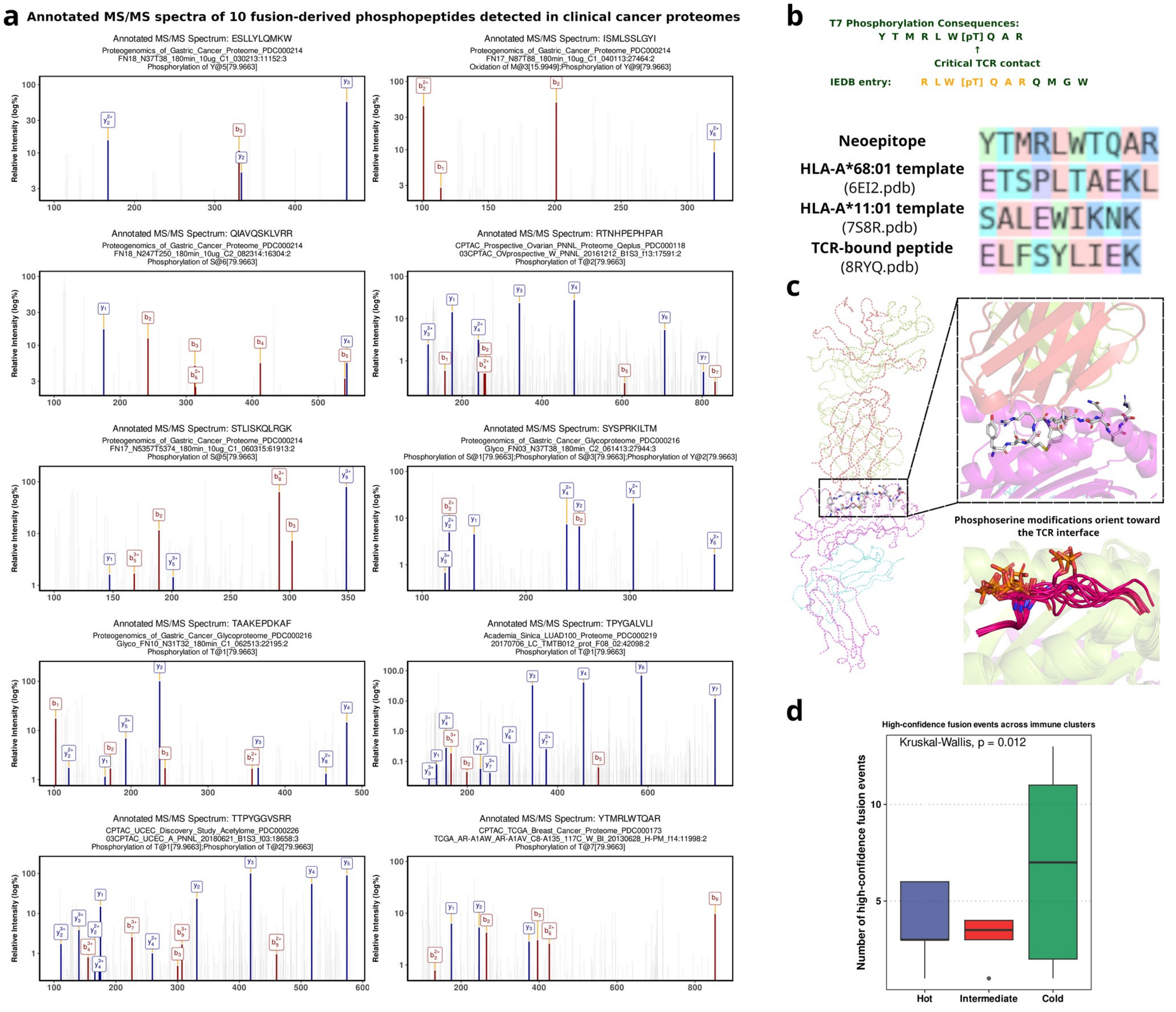


**Figure S7. Fusion-derived phosphopeptides as potential neoantigens and their clinical distribution.**

(a) Annotated MS/MS spectra of 10 fusion-derived phosphopeptides detected in clinical cancer proteomes. ESLLYLQMKW from ROCK2--TCF7L1 fusion with phosphorylation at Y5 (PDC000214); ISMLSSLGYI from BRD9--ABCA3 fusion with oxidation at M3 and phosphorylation at Y9 (PDC000214); QIAVQSKLVRR from TCF7L2--VTI1A fusion with phosphorylation at S6 (PDC000214); RTNHPEPHPAR from TCF7L2--VTI1A fusion with phosphorylation at T2 (PDC000118); STLISKQLRGK from DICER1--CLMN fusion with phosphorylation at S5 (PDC000214); SYSPRKILTM from SRSF10--MRPS21 fusion with phosphorylation at S1, S3, and Y2 (PDC000216); TAAKEPDKAF from CCDC97--BLVRB fusion with phosphorylation at T1 (PDC000216); TPYGALVLI from SH3GLB2--PHYHD1 fusion with phosphorylation at T1

(PDC000219); TTPYGGVSRR from SH3GLB2--PHYHD1 fusion with phosphorylation at T1 and T2 (PDC000226); YTMRLWTQAR from NISCH--PYROXD1 fusion with phosphorylation at T7 (PDC000173). Each panel shows the annotated MS/MS spectrum with b-ions (red) and y-ions (blue) labeled, demonstrating confident identification of fusion-derived phosphopeptides across multiple cancer proteomic datasets.

(b) Sequence alignment of the YTMRLWTQAR neoepitope. The query peptide was aligned with its variant (RLWTQARQMGW) and the peptide templates from the primary structural models (ETSPLTAEKL from 6EI2, SALEWIKNK from 7S8R, and ELFSYLIEK from 8RYQ). Alignment was performed using Clustal Omega based on the BLOSUM62 substitution matrix.

(c) Modelled YTMRLWTQAR-MHC-TCR complex. In all available crystal structures, phosphoserine residues are positioned on the TCR-facing surface of the pMHC complex (PDB ID: 3BGM, 3BH8, 3BH9, 3BHB, 3FQR, 3FQU, 3FQX, 3L6F, 4NNX, 4NO2, 4NO3, 5IEH, 7CIR, 7CIS, 7DYN).

(d) Distribution of high-confidence fusion events across immune clusters in the NCI–ICI cohort. High-confidence fusions were defined as coding fusion events jointly detected by STAR-Fusion and Arriba after filtering out immunoglobulin, mitochondrial, and same-gene artifacts, requiring FFPM > 0.1 (STAR-Fusion) or ≥2 split reads (Arriba), supported by ≥5 junction reads and at least one spanning fragment, and annotated as *YES_LDAS* with canonical or near-canonical splice junctions. Cold tumors exhibited a significantly higher number of fusion events compared to hot tumors ($p < 0.05$, Kruskal–Wallis test).
